# Accurate angular integration with only a handful of neurons

**DOI:** 10.1101/2022.05.23.493052

**Authors:** Marcella Noorman, Brad K Hulse, Vivek Jayaraman, Sandro Romani, Ann M Hermundstad

## Abstract

To flexibly navigate, many animals rely on internal spatial representations that persist when the animal is standing still in darkness, and update accurately by integrating the animal’s movements in the absence of localizing sensory cues. Theories of mammalian head direction cells have proposed that these dynamics can be realized in a special class of networks that maintain a localized bump of activity via structured recurrent connectivity, and that shift this bump of activity via angular velocity input. Although there are many different variants of these so-called ring attractor networks, they all rely on large numbers of neurons to generate representations that persist in the absence of input and accurately integrate angular velocity input. Surprisingly, in the fly, *Drosophila melanogaster*, a head direction representation is maintained by a much smaller number of neurons whose dynamics and connectivity resemble those of a ring attractor network. These findings challenge our understanding of ring attractors and their putative implementation in neural circuits. Here, we analyzed failures of angular velocity integration that emerge in small attractor networks with only a few computational units. Motivated by the peak performance of the fly head direction system in darkness, we mathematically derived conditions under which small networks, even with as few as 4 neurons, achieve the performance of much larger networks. The resulting description reveals that by appropriately tuning the network connectivity, the network can maintain persistent representations over the continuum of head directions, and it can accurately integrate angular velocity inputs. We then analytically determined how performance degrades as the connectivity deviates from this optimally-tuned setting, and we find a trade-off between network size and the tuning precision needed to achieve persistence and accurate integration. This work shows how even small networks can accurately track an animal’s movements to guide navigation, and it informs our understanding of the functional capabilities of discrete systems more broadly.

## INTRODUCTION

The brain is thought to rely on persistent internal representations of continuous variables for a wide range of computations, from working memory [1–3] to navigation [4–6] to motor control [7–9]. Such internal representations have been described in terms of low-dimensional manifolds along which higher-dimensional population activity evolves (Fig 1a), and they have been studied theoretically within the framework of continuous attractor networks [2, 10–17]; see [18–20] for recent reviews. One prominent example is the ring attractor network, a special class of continuous attractor networks that can maintain an internal representation of an angular variable, such as the orientation of self or of external stimuli [12, 21], and that has been proposed as a model of the head direction (HD) system [15, 22–31]. The underlying framework for ring attractor networks, which specifies the conditions under which a recurrently connected population of neurons can encode and update a continuous, angular variable, has historically relied on large numbers of neurons to ensure that these internal representations are both continuous and accurate. Here, we ask whether these same properties can be obtained in much smaller networks.

**Figure 1:**
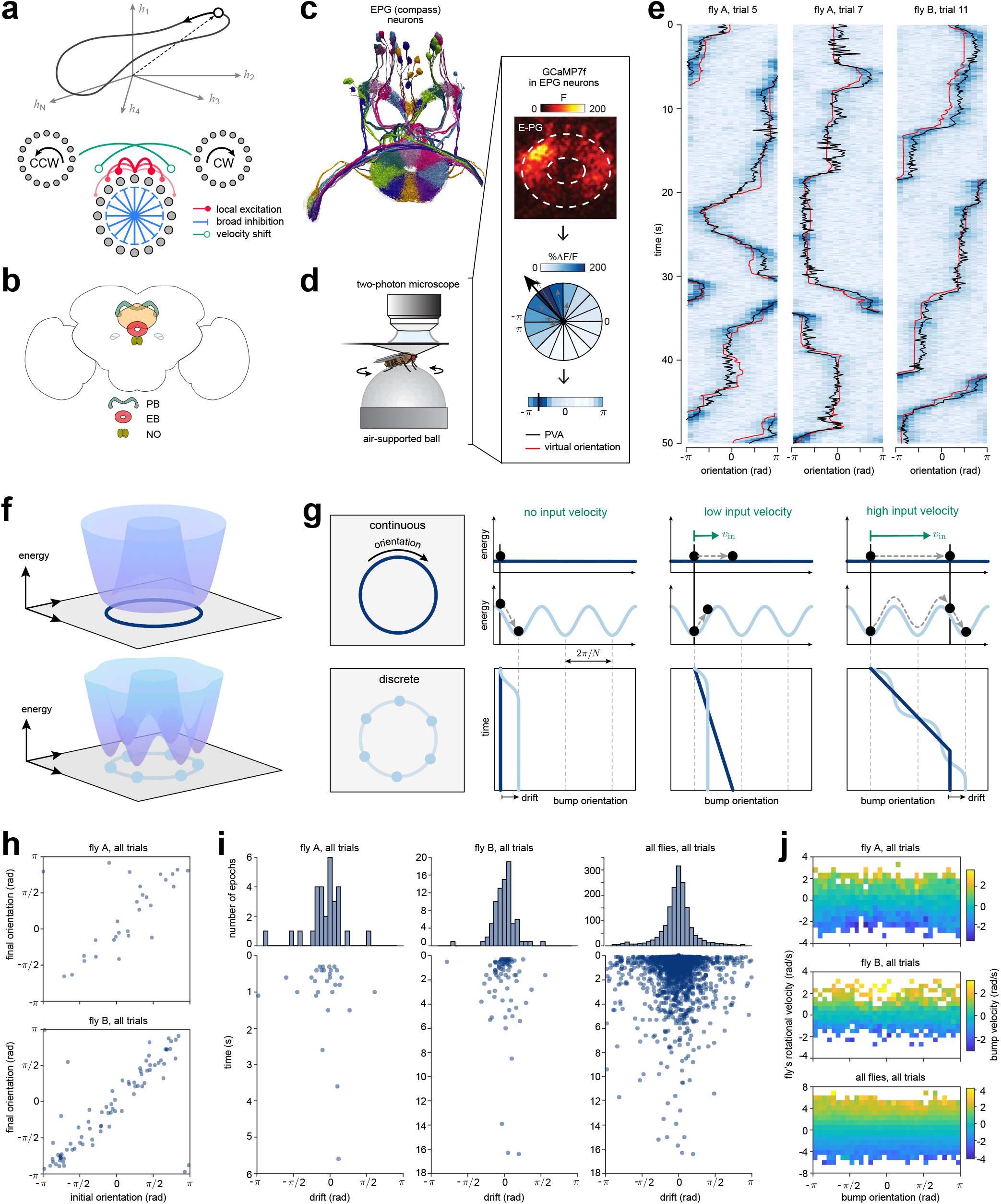
A biological attractor network can overcome hypothesized limitations of discreteness. **a)** Upper: Illustration of high dimensional neural activity that lies on a low dimensional, ring-like manifold. Lower: A ring attractor network can maintain an internal representation of orientation through a combination of local excitation (red) and broad inhibition (blue). Two side rings can update the internal representation based on angular velocity input through shifted connections back to the original ring (green). **b)** Schematic of the fly central complex. A population of compass neurons that innervate the ellipsoid body (EB) maintains an internal representation of the fly’s angular orientation as a localized bump of activity. A second population of shift neurons that innervate the protocerebral bridge (PB) shift the bump based on the fly’s angular velocity. The angular velocity input reaches the shift neurons through the noduli (NO). **c)** Electron microscopy reconstruction of compass neurons that are hypothesized to implement a ring-like attractor network. **d)** Two-photon imaging setup for tethered walking flies. Box: The ellipsoid body is segmented into 16 ROIs and used to compute the population vector average of the change in fluorescence Δ*F/F* (see *Methods* | *Data Analysis* | *Extracting Bump Orientation and Strength*). **e)** Compass neurons in the EB maintain a localized bump of activity (heatmap; population vector average in black) that tracks the fly’s angular orientation over time (red line). Shown for three different trials in two flies. **f)** In the absence of input, the dynamics of a ring-like attractor network will evolve toward the stable minimum of an energy landscape. Upper: Infinitely large networks generate energy landscapes with a continuum of marginally stable minima that lie along a ring. Lower: Small networks generate a bumpy energy landscape in which discrete energy minima are separated by barriers. **g)** The properties of the energy landscape lead to different dynamics in continuous (dark blue) versus discrete (light blue) networks. In continuous networks, the bump of activity will persist at the same orientation in the absence of input (second column) and will linearly integrate small and large velocities (third and fourth columns). In contrast, the discrete networks generate local energy minima that will attract the bump in the absence of input (second column), prevent the network from continuously integrating small velocity inputs (third column), and cause the network to nonlinearly integrate larger inputs (fourth column). **h)** Bump orientations in the EB before and after periods of stopping that exceeded 300 ms, shown for the same two flies as in panel **(e). i)** Distribution of bump drifts (top histogram), accumulated across periods of stopping from 300 ms to several seconds (lower panel). Shown for the same two flies (left and middle columns) and accumulated across 27 flies (right column). **j)** Average bump velocity (colormap) as a function of the fly’s rotational velocity for different bump orientations in the EB. Shown for the same two flies (upper two panels), and across all flies (lower panel). Gains between bump velocity and the fly’s rotational velocity on the ball in darkness tend to vary by individual; average bump velocities shown across all flies (lower panel) were normalized for these gain differences. See *Methods* |*Data Analysis* |*Characterizing Bump Drift* and *Methods* |*Data Analysis* |*Characterizing Bump Velocity* for details of analysis.

Ring attractor networks get their name from the one-dimensional ring manifold on which activity evolves. This manifold emerges in the limit that an infinitely large population of HD-tuned neurons maintains sustained and localized activity via positive feedback; one way that this can be achieved is through recurrent connectivity by which neurons with similar tuning excite one another, and neurons with dissimilar tuning inhibit one another (Fig 1a, [12, 21, 23, 32], but also see [33]). The resulting population dynamics can generate a localized bump of activity that persists at the same orientation in the absence of input and traverses the ring manifold through the integration of self-motion inputs [22, 23, 28]. As a result of their infinite size, ring attractor networks achieve infinite precision in maintaining and accurately updating the bump of activity.

Although ring attractor networks were first proposed by theorists several decades ago, it has been difficult to identify actual ring-like network architectures in brains. Notably, ring attractor networks have been invoked to explain experimental observations of bell-shaped tuning curves of mammalian HD cells that display persistent firing in the absence of input and whose activity is updated by self-motion even in darkness [4, 34], but it has not yet been possible to measure the patterns of connectivity between these cells. Mammalian HD neurons have been observed to change their tuning coherently when animals are placed in different settings [4], and recent work suggests that the HD population dynamics do traverse a one-dimensional ring-like manifold [35]. In the fly *Drosophila melanogaster*, a recurrent network of neurons in a brain region called the central complex (CX, Fig 1b) was recently shown to exhibit the functional and structural connectivity (Fig 1c, [36–38]), and the dynamics (Fig 1d-e, [6, 36, 39, 40]), of a ring-like attractor network. These dynamics are observable as a bump of population activity in so-called EPG or “compass” neurons in a toroidal structure of the CX called the ellipsoid body (EB). This bump of activity tracks the fly’s orientation as the fly turns and maintains its orientation during epochs when the fly stops moving (Fig 1e). These dynamics are driven both by localizing sensory cues and by the integration of self-motion cues, which enables the bump to track the fly’s movements even in darkness [6, 39, 40]. The architecture of the circuit features two subpopulations of “shift” neurons that are jointly tuned to orientation and angular velocity and that receive input from and project back to the compass neurons [37–40], as previously hypothesized in theoretical models (Fig 1a) [22]. Thus, both physiological and anatomical considerations suggest that this circuit exhibits the key features of a ring-like attractor network, with one major exception: the fly circuit has far fewer computational units—sets of neurons with the same HD tuning—than are thought necessary to approximate an accurate ring attractor [38]. In what follows, we dissect the functional properties of discrete ring-like attractor networks, and we show how small circuits might overcome the limitations of this discreteness to achieve functional performance thought to emerge only in the limit of large systems.

## RESULTS

The computational properties that make ring attractor networks such appealing models of the HD system arise in the limit of large system sizes. Specifically, in the limit as the number of neurons approaches infinity (what we will refer to as a “continuous” system), a ring attractor network generates a continuum of configurations that define the ring attractor manifold (Fig 1f, upper panel). These configurations are marginally stable, such that perturbations along the ring attractor manifold will be maintained, whereas perturbations off the manifold will result in activity that quickly returns to it. These properties allow us to express the manifold as a flat dimension in the energy landscape of the system; all points along this flat dimension have equal and minimum energy, and thus the system can stably sit at any of these points in the absence of input (Fig 1g, second column, dark blue). Moreover, small changes in input can drive the system along this flat dimension without obstruction, such that the population activity accurately tracks these changes (Fig 1g, third/fourth columns, dark blue). This flat energy dimension gives the system infinite precision in encoding and updating an internal representation of a one-dimensional circular variable such as HD.

However, when the system is small (what we will refer to as a “discrete” system), these properties break down, thereby limiting how precisely the internal HD representation can be stored and updated. Instead of exhibiting a flat dimension, the energy landscape exhibits a set of discrete basins (Fig 1f, lower panel) that attract the population activity in the absence of input (Fig 1g, second column, light blue), prevent the integration of small inputs (Fig 1g, third column, light blue), and prevent the accurate integration of large inputs (Fig 1g, fourth column, light blue). For a small network like the fly compass network, we would thus expect to observe three distinct signatures of discreteness: (1) drift in the absence of input, in which the HD bump drifts to stereotyped orientations around the EB when the fly stops turning; (2) failure to integrate small angular velocities, in which the HD bump does not move continuously when the fly makes small turns; and (3) variable responses to larger angular velocities, in which the HD bump moves faster or slower relative to the fly’s movements, depending on its orientation within the EB.

To assess whether the fly circuit is capable of overcoming these expected shortcomings of discreteness, we analyzed the dynamics of the HD bump by performing two-photon calcium imaging of the compass neurons in the EB while head-fixed flies walked on an air-supported ball in darkness (Fig 1d,e,h-j; see *Methods* |*Experimental Setup* and *Methods* |*Data Analysis*). While there is fly-to-fly variability in the accuracy of angular integration, which may be due in part to limitations of the fly-on-a-ball system (see *Methods* |*Experimental Setup* |*Spherical Treadmill System*), several flies showed a remarkable ability to track changes in their angular orientation in darkness. We first measured how far the HD bump would drift in the absence of input [6]. To do this, we compared the orientation of the HD bump in the EB when the fly stopped moving to when the fly began walking again; we did not observe any apparent signatures that the bump drifted to a discrete number of stereotypical orientations in the EB (Fig 1h), as might be expected from the small number of computational units involved. The distribution of drifts was strongly peaked at zero (Fig 1i, upper row), and included epochs in which the bump persisted at the same orientation for several seconds (Fig 1i, lower row) [6]. We then analyzed the average bump velocity as a function of the fly’s average turning velocity when the bump was in different positions in the EB. Again, across several flies, we saw that the bump velocity was consistent across different orientations in the EB, with no apparent signatures of nonlinear integration nor any indication that the bump failed to track small velocities (Fig 1j). Thus, despite the imperfections of measuring the HD system’s performance in head-fixed flies walking on a ball, we found that the peak performance of the HD system belied its small size in both its low drift and in the accuracy of its integration of angular velocity.

### Optimally-tuned discrete networks generate a continuum of stable bump configurations

The previous results suggest that small networks can in practice integrate angular velocity without suffering the performance failures expected of discrete systems. To explore how such accurate integration might be achieved in principle, we studied the performance of small attractor networks (Fig 2a; see *Methods* |*Model Overview*). We considered networks of *N* heading-tuned neurons whose preferred headings *θ*_*j*_ uniformly tile heading space, with an angular separation of Δ*θ* = 2π*/N* radians. These neurons can be arranged topologically in a ring according to their preferred headings, with neurons locally exciting and broadly inhibiting their neighbors. We capture this with a symmetric cosine weight matrix 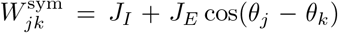, where *J*_*E*_ and *J*_*I*_ respectively control the strength of the tuned and untuned components of recurrent connectivity between neurons with preferred headings *θ*_*j*_ and *θ*_*k*_. We will refer to these components as local excitation and broad inhibition, respectively (but note that the tuned component takes on both positive and negative values, and thus is not strictly excitatory; within the parameter regimes that we will consider, the untuned component will be strictly inhibitory). The network receives angular velocity input *v*_in_ through asymmetric, velocity-modulated weights 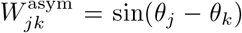 (see also [23]); this input could be implemented via two linear side rings whose time constants are much smaller than the time constant of neurons in the center ring (see *Supplemental Information* |*Network Equations* for more details). We further assume that each neuron transforms its inputs via a nonlinear transfer function *ϕ* (·). The total input activity *h*_*j*_ of each neuron is governed by:

**Figure 2:**
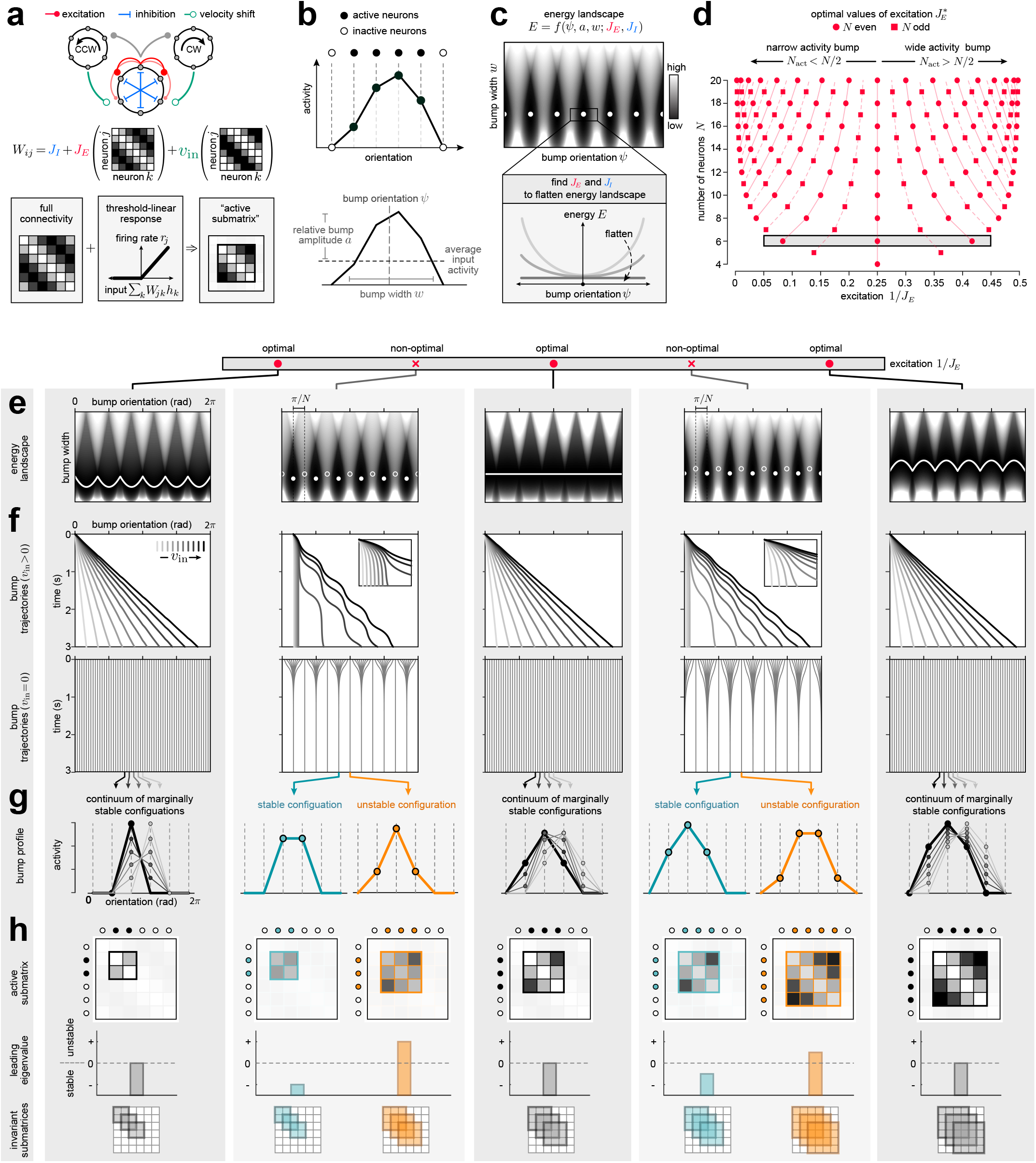
Optimally-tuned local excitation can recover a ring attractor manifold. **a)** Schematic of network model. Upper: A population of neurons is recurrently connected via local excitation of strength *J*_*E*_ and broad inhibition of strength *J*_*I*_. Two side rings receive input from the center ring and project back to the ring with shifted, velocity dependent connections. Lower: Neurons transform their input through a threshold linear response function; within the parameter regime considered here, this ensures that only a subset of neurons are active at any time. The dynamics of this subset is governed by an “active submatrix” of the full connectivity. **b)** Upper: The strength of local excitation and broad inhibition can be tuned to achieve a bump of population activity that is maintained by a subset of *N*_act_ active neurons (filled circles). Lower: The configuration of this bump can be specified by its width *w*, amplitude relative to average input activity *a*, and orientation *ψ*. Note that the average input activity (black dashed line), is incorporated into the bump width *w*, and is thus unnecessary to include in the bump configuration (see *Supplemental Information* |*Network Equations* |*Order Equations* for more details). **c)** Upper: For a given set of parameters {*J*_*E*_, *J*_*I*_}, we can specify the energy of different bump configurations. When these parameters are naively tuned, the resulting energy landscape is bumpy, with local minima (white points) separated by barriers. Lower: To optimize network performance, we searched for the parameters that minimize the local curvature as a function of bump orientation and thereby “flatten” the energy landscape. **d)** For a given network size *N*, we find that there are *N* 3 values of local excitation that will flatten the energy as a function of bump orientation. These can be organized by whether the bump is maintained by greater (*N*_act_ *> N/*2) or fewer (*N*_act_ *< N/*2) than half the neurons in the network. Shaded bar indicates these values of local excitation for a network size of *N* = 6, expanded upon in panels **(e-h). e-h)** We evaluate network performance (rows) for networks of size *N* = 6 and for different values of local excitation (columns); we consider all three optimal values 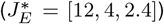, and two intermediate values (*J*_*E*_ = [6, 3]). **e)** Optimal networks generate energy landscapes that are flat as a function of orientation (white line), while non-optimal networks generate landscapes that have local minima (filled markers) separated by barriers (open markers). **f)** Optimal networks can linearly integrate angular velocity inputs (upper row), and the bump persists at the same orientation in the absence of input (lower row). Non-optimal networks fail to integrate small inputs (upper row, insets) and nonlinearly integrate larger ones (upper row, main panels); when this input is removed, the bump will drift to the nearest local minimum of the energy landscape (lower row). **g)** In optimal networks, the bump of activity can persist at a continuum of different orientations in the absence of input. Non-optimal networks can persistently maintain a discrete number of bump configurations; one set of these is stable (turquoise; corresponding to filled markers in **(e)**), and the other is unstable (orange; corresponding to open markers in **(e)**). See *Methods Model Simulations* |*Drift in the Absence of Input* and *Methods* |*Model Simulations* |*Velocity-Driven Dynamics* for simulation details. **h)** In all networks, the persistent bump of activity in **(g)** is maintained by a subset of active neurons (filled circles) whose dynamics are governed by the leading eigenvalue of an active submatrix of the full connectivity (top row, see also panel **(a)**). In optimal networks, the bump is always maintained by the same number of active neurons, and thus these active submatrices always have the same size (bottom row). The leading eigenvalue of these submatrices is zero (middle row), indicating that each active submatrix generates a line attractor [10]. In non-optimal networks, the bump is maintained by different numbers of active neurons depending on whether the bump configuration is stable (turquoise) or unstable (orange) (bottom row). The corresponding active submatrices differ in size and have eigenvalues that are less than or greater than zero (middle row), indicating that these submatrices generate stable and unstable fixed points, respectively.

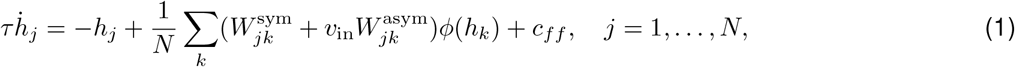

where *c*_*ff*_ is a constant feedforward input to all neurons in the network. In what follows, we will take *ϕ* (·) to be threshold linear; for the parameter regimes that we will study here, this ensures that only a strict subset of all neurons will be active at any time. As a result, the dynamics of these active neurons will be governed by an “active submatrix” of the full connectivity (Fig 2a, bottom).

For sufficiently strong local excitation and broad inhibition, this network will generate a stable bump of activity (Fig 2b, upper panel; see SI Fig S1 and *Methods* |*Model Analytics* |*Stationary Solutions*); we will work within this stable parameter regime for the remainder of our analyses. We describe this bump of activity via the Fourier modes of the population activity (given by Eq. 1). For the network connectivity chosen here, which varies sinusoidally as a function of the difference between preferred headings, the population activity can be fully specified by the zeroeth- and first-order Fourier modes. This allows us to fully characterize the “configuration” of the activity bump in terms of its relative amplitude *a*, angular width *w*, and angular orientation *ψ* (Fig 2b, lower panel; see also *Supplemental Information* |*Network Equations* |*Order Equations*). These quantities, which are computable from the Fourier modes, vary continuously over time, and thus the same number of active neurons can maintain bump configurations with a range of different relative amplitudes, widths, and orientations.

We began by characterizing the manifold of stable bump configurations in the absence of angular velocity input. To this end, we constructed a landscape that describes the energy of different bump configurations for a given set of parameters *J*_*E*_ and *J*_*I*_ [41, 42] (see *Methods* |*Model Analytics* |*Energy Landscape*). For the majority of parameter settings, the energy landscape was bumpy, with discrete minima separated by energy barriers (Fig 2c, upper panel). The energy landscape about these minima exhibited high local curvature as a function of orientation, indicating that the bump would be highly attracted to these particular orientations. To weaken this attraction, we analytically determined the values of *J*_*E*_ and *J*_*I*_ that would locally minimize this curvature, and thus locally flatten the energy landscape (Fig 2c, lower panel). Surprisingly, we found that specific values of local excitation could drive the curvature to zero, resulting in an energy landscape that was completely flat as a function of orientation (SI Fig S2). For a network of size *N*, there are *N* − 3 such “optimal” values of local excitation 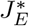(Fig 2d). Fig 2e illustrates the corresponding optimal energy landscapes for a network of size *N* = 6, and contrasts these landscapes with two non-optimal landscapes generated using intermediate values of local excitation.

To verify that these optimally-tuned networks could overcome the failure modes highlighted in Fig 1g, we simulated the response of each network to a constant velocity input (Fig 2f; see *Methods* |*Model Simulations* |*Overview*). As expected, we found that optimal networks could accurately integrate a range of different input velocities, such that the orientation of the bump changed linearly over time (Fig 2f, upper row). When this velocity input was removed (Fig 2f, lower row), the bump persisted at the same orientation and did not drift (note that even in networks with different nonlinearities and connectivity profiles, we find parameter values for which this is the case; see SI Fig S3 and *Methods* |*Model Simulations* |*Robustness to Changes in the Transfer Function and Recurrent Weights*). In contrast, non-optimal networks failed to integrate small velocities (Fig 2f, upper row insets), and they nonlinearly integrated larger velocities (Fig 2f, upper row main panels). When this velocity input was removed, the bump drifted toward the set of discrete orientations that correspond to local minima of their energy landscapes (Fig 2f, lower row).

In the absence of velocity input, the optimal networks generate a continuum of marginally stable configurations in which the bump can persist (Fig 2g). These configurations share one striking feature: the bump is always maintained by the same number of active neurons, despite variations in the relative amplitude, width, and orientation of the bump. This feature has important consequences for the dynamics of the network: when a fixed subset of neurons is active, the corresponding equations (Eq. 1 for *h*_*j*_ *>* 0) reduce to a linear dynamical system that depends only on an “active submatrix” of the full connectivity *W* (Fig 2h, upper row; note that here, we take the full connectivity to be *W* = (*W* ^sym^*/N* − *I*)/ τ). Moreover, because the connectivity is rotationally invariant, this active submatrix—and thus the resulting network dynamics—will be identical for any contiguous subset of *N*_act_ active neurons. To characterize these dynamics, we determined the eigenvalue spectra of these active submatrices (see *Methods* |*Model Analytics* |*Leading Eigenvalues of Active Submatrices*). We found that each exhibited a single zero eigenvalue (Fig 2h, middle row), with the real part of all remaining eigenvalues less than zero. This property gives rise to a so-called line attractor that produces a continuum of marginally stable configurations along a line [10]. Thus, in this network, a ring attractor emerges as a discrete set of *N* line attractors that govern the dynamics of distinct subsets of active neurons (Fig 2h, lower row), and that are “stitched together” at the points where an active subset gains and loses an active neuron.

In contrast, the non-optimal networks can only maintain a discrete set of bump configurations in the absence of input; these configurations correspond to so-called fixed points of the dynamics. One subset of these configurations is stable, and the bump will return to these configurations following small perturbations (Fig 2g, turquoise curves).

The other subset is unstable, and the bump will move away from these configurations if perturbed (Fig 2g, orange curves). In these two configurations—stable and unstable—the bump is maintained by different numbers of active neurons (also called the “support” of the fixed point; [43, 44]), and the corresponding active submatrices differ in size (Fig 2h, upper row). The smaller of these submatrices has a leading eigenvalue less than zero and governs network dynamics about the stable fixed point, while the larger of these submatrices has a leading eigenvalue greater than zero and governs dynamics about the unstable fixed point (Fig 2h, middle row). In what follows, we use these active submatrices to dissect the dynamics of non-optimal networks in a manner that is not easily extractable from the energy landscape itself, and we show how the balance between stable and unstable dynamics shapes network performance.

### Non-optimal networks vary in how they balance periods of stability and instability

The previous results highlight a unique feature of threshold linear networks: when a fixed subset of neurons is active, the corresponding dynamical system is linear, and the dynamics of the full network can be viewed as a set of linear subsystems (Fig 3a, upper panel). The recurrent connectivity assumed here will ensure that each of these subsystems is defined by a contiguous subset of neurons that actively maintain the heading bump (Fig 3a, middle panel). Within each linear subsystem, the dynamics of the active neurons (and hence the orientation of the bump) are entirely determined by the eigenvalues of the active submatrix of *W*. Specifically, the activity of each neuron will be given by a sum of exponential functions whose rates are given by these eigenvalues; assuming a sufficiently large gap between the leading eigenvalue *λ* and all other eigenvalues, the dynamics of each active neuron will be dominated by the exponential with rate *λ*. It follows that the orientation of the bump will also evolve exponentially over time with a rate *λ* (Fig 3a, lower panel):

**Figure 3:**
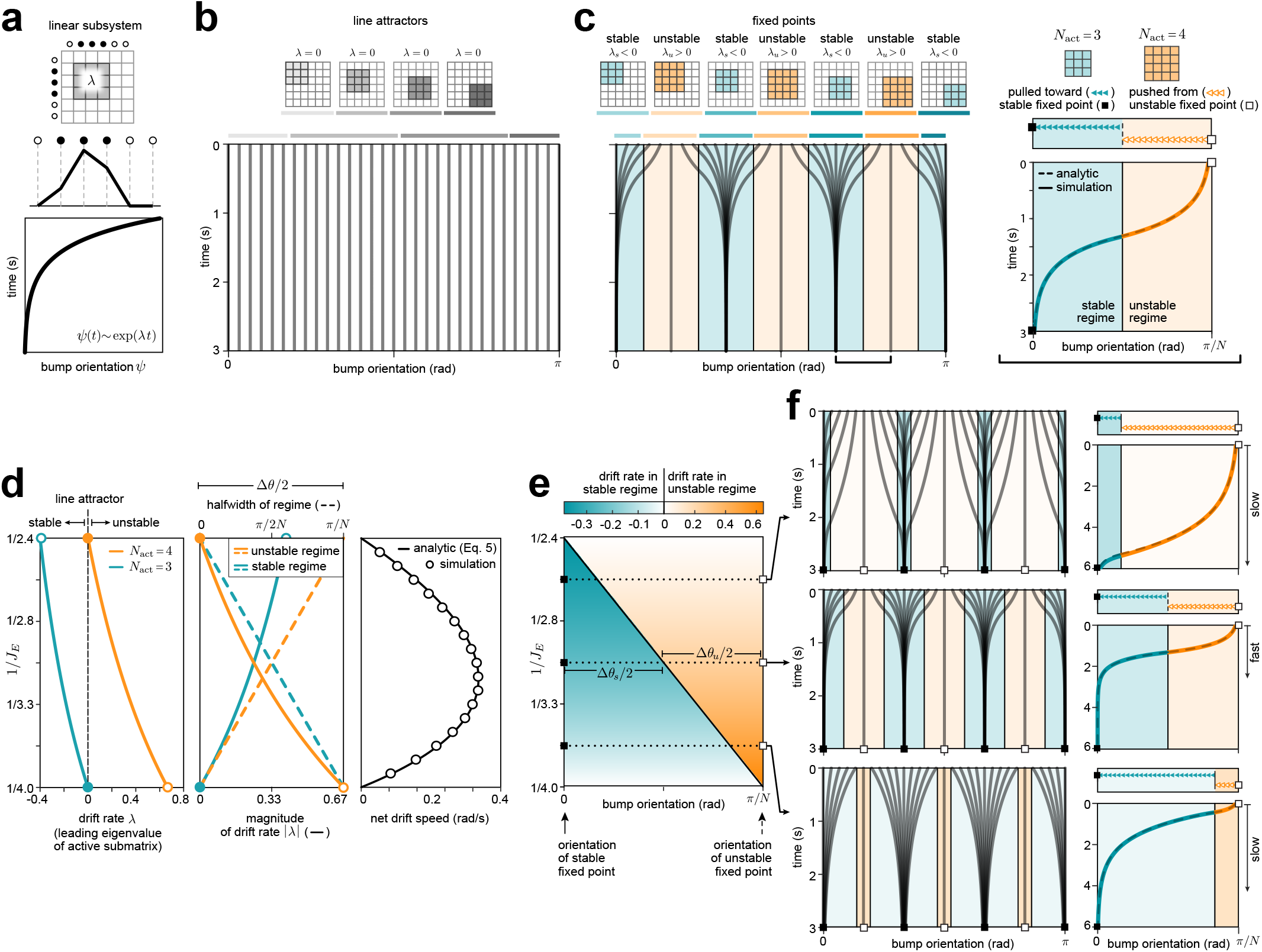
Non-optimal networks exhibit periods of stability and instability that give rise to drift in the absence of input. **a)** When a fixed subset of neurons is active, the network dynamics are linear and are governed by an active submatrix of the connectivity with leading eigenvalue *λ* (upper). These dynamics will generate a bump of activity (middle) whose orientation changes exponentially over time with a rate *λ* (lower). **b)** In optimal networks, the bump is maintained by a fixed number of active neurons (shown for *N*_act_ = 3, corresponding to 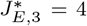). The corresponding active submatrices have leading eigenvalues of zero, and thus the bump does not drift over time, regardless of its initial orientation. See *Methods* |*Model Simulations* |*Drift in the Absence of Input* for simulation details. **c)** Left: In non-optimal networks, the bump is maintained by two different numbers of active neurons (shown for *J*_*E*_ = 3). One of the corresponding active submatrices has an eigenvalue that is greater than zero, and generates unstable dynamics in which the bump is pushed from an unstable fixed point (“unstable regime”, orange; *N*_act_ = 4); the other has an eigenvalue less than zero, and generates stable dynamics in which the bump is pulled toward a stable fixed point (“stable regime”, turquoise; *N*_act_ = 3). Over time, the bump will drift away from its initial orientation and toward the stable fixed point. Right: illustration of single trajectory in which the bump drifts from the unstable regime to the stable one, computed analytically (dashed line) and via simulation (solid line). **d)** Left: Optimal values of local excitation generate a single active submatrix with a leading eigenvalue of zero (filled circles, shown for 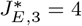 (turquoise) and 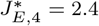(orange), corresponding to *N*_act_ = 3 and *N*_act_ = 4, respectively). As the local excitation is weakened away from the larger of these optimal values, the leading eigenvalue of the smaller active submatrix decreases below zero (turquoise); this eigenvalue defines the (negative) drift rate *λ*_*s*_ of the stable regime. Conversely, as the local excitation is strengthened away from the smaller of these optimal values, the leading eigenvalue of the larger active submatrix increases above zero (orange); this eigenvalue defines the (positive) drift rate *λ*_*u*_ of the unstable regime. Middle: Changes in the magnitude of drift rates |*λ*_*s*_| and |*λ*_*u*_| (solid lines) are offset by changes in the halfwidths of the stable and unstable regimes (dashed lines). At one extreme, as the magnitude of the stable drift rate approaches zero (solid turquoise line), the halfwidth of the stable regime approaches *θ/*2 = π*/N*, and the halfwidth of the unstable regime approaches zero (dashed lines). At the other extreme, the reverse is true; as the magnitude of the unstable drift rate approaches zero (solid orange line), the halfwidth of the unstable regime approaches *θ/*2 = π*/N*, and the halfwidth of the stable regime approaches zero (dashed lines) Right: The net drift speed will be largest (fastest drift) when the local excitation is intermediate between two optimal values, and will shrink to zero (no drift) as the local excitation approaches an optimal value. To measure this speed, we compute the time *τ*_*d*_ that it takes to drift a fixed angular distance *Δψ*_*d*_, and we use Δ ψ_*d*_/τ_*d*_ as a measure of the net drift speed (see *Methods* |*Model Analytics Drift in the Absence of Input*). **e)** The bump dynamics are governed by the orientations of the stable and unstable fixed points (filled and open squares, respectively), the halfwidths Δ*θ*_*s*_/2 and Δ*θ*_*u*_/2 of the stable and unstable regimes (turquoise and orange regions, respectively), and the drift rates within each regime (color saturation). Note that, consistent with the middle panel of **(d)**, increases in the magnitude of the drift rate (deeper color saturation) are accompanied by decreases in the width of the regime over which that drift rate applies. **f)** Different values of local excitation produce qualitatively different drift trajectories, shown for many different initial conditions (left column) and dissected for one initial condition in which the bump begins near an unstable fixed point (right column). In the unstable regime (orange), the bump accelerates away from an unstable fixed point, and the trajectory has positive curvature; in the stable regime (turquoise), the bump decelerates toward a stable fixed point, and the trajectory has negative curvature. The bump drifts more rapidly for values of local excitation that are far from optimal (middle panel), consistent with panel **(d)**. Right column is computed both analytically (dashed lines) and via simulation (solid lines).

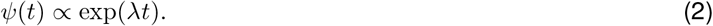

When this rate is positive, the bump will move exponentially quickly away from a single unstable fixed point; when this rate is negative, the bump will move exponentially slowly toward a single stable fixed point. In the special case that the rate is zero, the bump will not move. The fact that all optimal networks produce a set of zero eigenvalues thus ensures that the resulting network dynamics will not drift in the absence of input (Fig 3b). In contrast, all non-optimal networks have two sets of eigenvalues—one positive, *λ*_*u*_, and one negative, *λ*_*s*_—that give rise to unstable and stable network dynamics, respectively (Fig 3c, left). These dynamics govern the behavior of the bump in the vicinity of an unstable or stable fixed point (note that in the absence of input, these fixed points are spaced by Δ*θ/*2 = π*/N* radians; see Fig 2e and *Methods* |*Model Analytics* |*Stationary Solutions*). In the absence of velocity input, the bump will drift over time from an unstable regime to a stable regime, which will be accompanied by a decrease in the number of neurons that actively maintain the bump (Fig 3c, right).

The value of local excitation in the network will determine the leading eigenvalues of the active submatrices, and thus the drift rates in the stable and unstable regimes. If we vary the local excitation between two optimal values 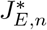 and 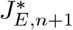 (corresponding to scenarios in which the bump is always maintained by *n* or *n* +1 active neurons, respectively), we find that the drift rates vary depending on how closely tuned the local excitation is to either optimal value (Fig 3d, left; SI Fig S4; *Methods* |*Model Analytics* |*Leading Eigenvalues of Active Submatrices*):

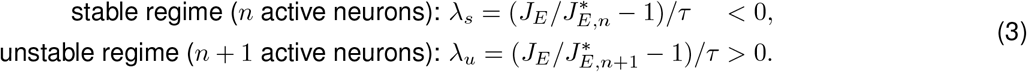

At one extreme, when the local excitation is close to 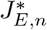 the drift rate in the *stable* regime goes to zero. At the other extreme, when the local excitation is close to 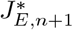, the drift rate in the *unstable* regime goes to zero. In between these extremes, there are moderate drift rates in both regimes.

Each stable and unstable regime governs the dynamics of the bump across a range of bump orientations. We use this range of orientations to define the angular widths Δ*θ*_*s*_ and Δ*θ*_*u*_ of each regime, which depend on the ratio of stable and unstable drift rates (see *Methods* |*Model Analytics* |*Widths of Stable and Unstable Regimes*):

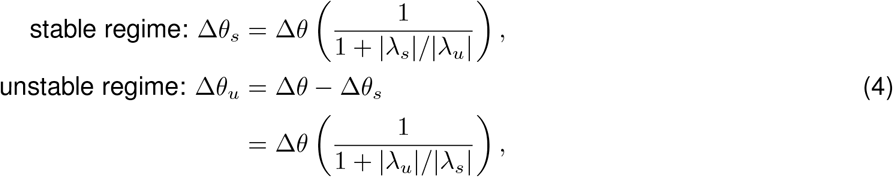

where Δ*θ* is the spacing between preferred headings. At one extreme, as _*s*_ approaches zero, the stable regime grows to fill the entire width Δ*θ*, and the unstable regime shrinks to zero. At the other extreme, as *λ*_*u*_ approaches zero, the unstable regime grows to fill the entire width Δ*θ*, and the stable regime shrinks to zero (Fig 3d, middle). The net drift speed | *λ*_*d*_|, which quantifies how quickly the bump moves across a fixed angular interval, can be expressed as a product between the width of either regime and the rate in that regime (see *Methods* |*Model Analytics* |*Drift in the Absence of Input*):

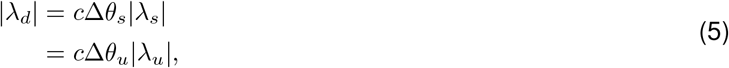

where *c* = (*e* − 1)/2*e* is a constant. As a result, increases in the width of each regime are offset by decreases in the rate within that regime (Fig 3d, middle), such that the net drift speed approaches zero as the local excitation approaches an optimal value (Fig 3d, right).

Together, this analysis enables us to dissect the bump dynamics in terms of three factors (Fig 3e): (i) the angular orientations of the stable and unstable fixed points, (ii) the angular widths of the stable and unstable regimes about these fixed points, and (iii) the drift rates in these stable and unstable regimes. Together, these three factors determine how quickly the bump will drift over time, and they determine the precise shape of the drift trajectory (Fig 3f). In the absence of velocity input, we can analytically determine all three factors from the symmetric connectivity *W* ^sym^. Velocity input will introduce an asymmetric component *v*_in_*W* ^asym^ to the connectivity (see Fig 2a); however, for sufficiently small velocities, this component will have a negligible effect on the angular widths and drift rates of the stable and unstable regimes (SI Fig S5; see *Methods* |*Model Analytics* |*Small Velocity Approximation* for details). This enables us to dissect the dynamics of velocity integration in terms of changes in the orientations of the stable and unstable fixed points, as we describe in the next section.

### The orientations of stable and unstable fixed points impact velocity integration

In the absence of velocity input, the bump can stably persist at one of *N* different stable fixed points. When a small and constant velocity input *v*_in_ is injected into the network, the orientation of each stable fixed point shifts in the direction of the velocity input (Fig 4a, solid lines), and the bump will be driven toward this new stable fixed point. For sufficiently small velocities, this fixed point will remain within the stable regime, and the bump will converge to it (turquoise trajectories in Fig 4b, “below threshold velocity” panels; see *Methods* |*Model Analytics* |*Locations of Fixed Points in Velocity-Driven Regime*).

**Figure 4:**
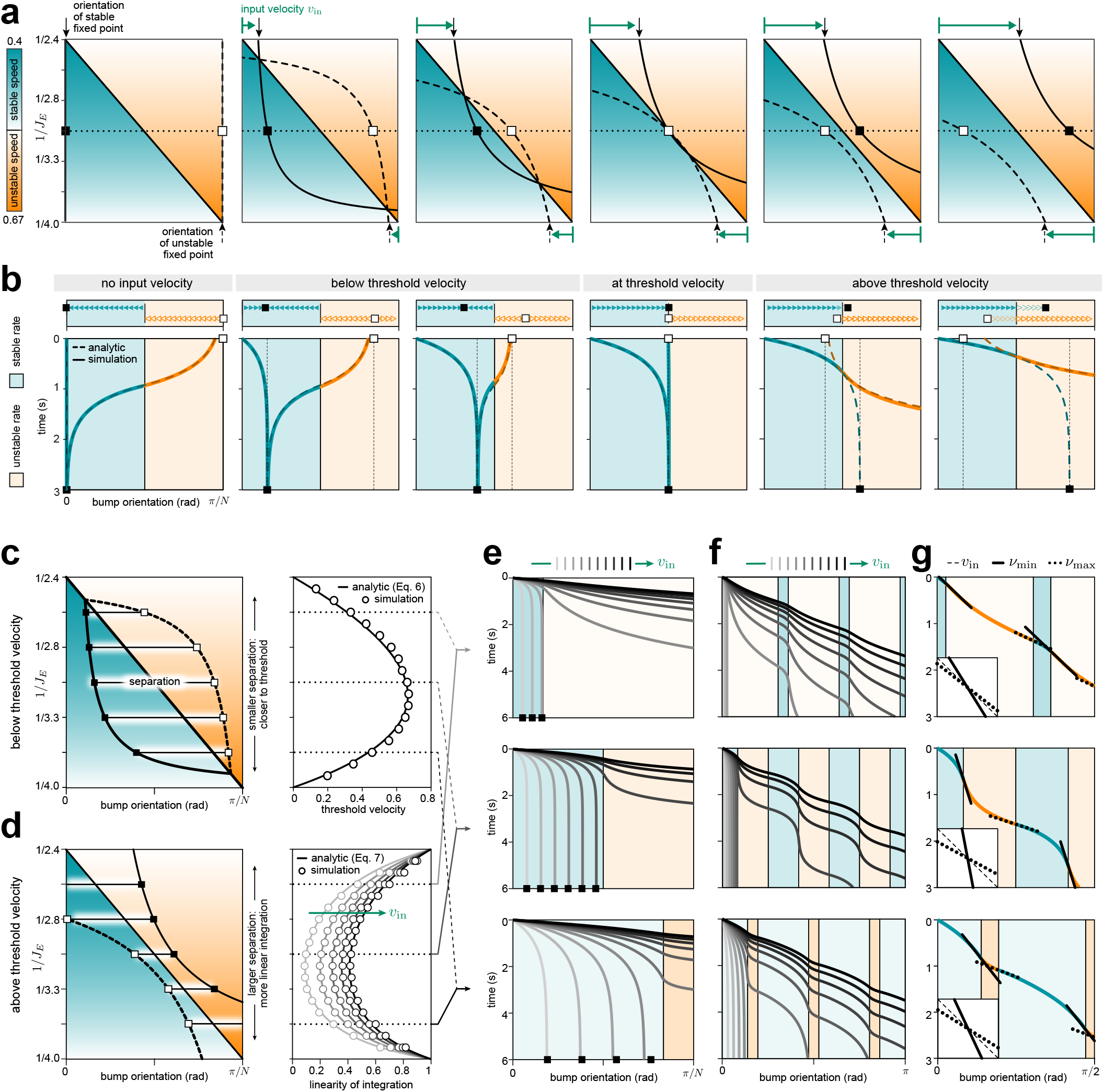
The orientations of stable and unstable fixed points govern how non-optimal networks integrate velocity inputs. **a)** Orientations of stable and unstable fixed points as a function of velocity input. Left panel: In the absence of velocity input, the orientations of stable and unstable fixed points do not depend on the value of local excitation. Middle and right panels: as velocity input increases (green arrows), the stable fixed points (solid lines) shift with the velocity, and the unstable fixed points (dashed lines) shift against the velocity, such that both approach and pass through the boundary between the stable and unstable regimes. Filled and open squares respectively mark the orientations of stable and unstable fixed points considered in panel **(b). b)** Bump trajectories for the same velocities as in panel **(a)**, but for a fixed value of local excitation *J*_*E*_ = 3. When the stable and unstable fixed points lie within their respective regimes, the bump will be pulled toward the stable fixed point (“below threshold velocity” panels); this can result in the bump moving in the opposite direction as the velocity input (orange/turquoise curves). At a particular threshold velocity, the two fixed points will meet at the boundary between regimes (“threshold velocity” panel). Above this velocity (“above threshold velocity” panels), the bump will have sufficient input to move out of the stable regime and into the unstable regime. The stable fixed point will move beyond the stable regime, and thus the bump will be pulled from ahead by a stable fixed point that it can never reach (solid and dashed turquoise lines; note that the solid line follows the dashed line toward the stable fixed point up until it reaches the boundary with the unstable regime. The continuation of the dashed line beyond the boundary indicates the trajectory that the bump would have taken if the number of active neurons did not change, determined analytically). Once the bump moves into the unstable regime, it will be pushed from behind by an unstable fixed point that is beyond the unstable regime (solid and dashed orange lines; again, note that the solid line begins only when the bump crosses into the unstable regime, but follows the dashed line that leads away from the unstable fixed point). Bump trajectories are computed analytically (dashed lines) and via simulation (solid lines). **c)** Left: Orientations of stable (solid lines) and unstable (dashed lines) fixed points as a function of local excitation for a single input velocity below threshold. The bigger the separation between stable and unstable fixed points (left), the larger the threshold velocity needed to move the bump continuously (right; determined analytically (solid lines) and via simulations (markers)). See *Methods* |*Model Simulations* |*Measuring Threshold Velocity*. **d)** Left: Same as left panel of **(c)**, but for a single input velocity above threshold. The smaller the separation between stable and unstable fixed points (left), the larger the effect the fixed points will have on the dynamics, and the more nonlinear the resulting velocity integration (right). We measure the linearity of integration as the ratio of the slowest and fastest bump speeds for a given constant velocity input; see *Methods Model Simulations Measuring Linearity of Integration*. **e, f)** Example bump trajectories that highlight behavior below threshold **(e)** and above threshold **(f)**, generated using the values of local excitation highlighted in panels **(c**,**d)**. Top and bottom rows: For values of local excitation that are closer to optimal, small input velocities will drive the bump past the edge of the stable regime, leading to continuous integration for a smaller input **(e)**. The resulting trajectories have local kinks that are more quickly flattened out as velocity increases **(f)**. Middle row: For values of local excitation that are far from optimal, the network requires higher input velocities to drive the bump past the edge of the stable regime **(e)**, and the resulting trajectories are more nonlinear **(f)** (consistent with **(c)** and **(d)**, respectively). **g)** For velocities above threshold, the bump will be moving the slowest as it leaves the stable regime (solid lines) and the fastest as it leaves the unstable regime (dotted lines). The difference in fastest and slowest bump velocities will be most pronounced for values of local excitation that are far from optimal (middle panel). See *Methods* |*Model Simulations* |*Parameter Choices* for specific input velocity values used in each panel.

The same velocity input will also shift the orientation of the unstable fixed points, but in the opposite direction (Fig 4a, dashed lines). As a result, if the initial bump has not yet settled into a stable fixed point when the velocity turns on, the bump can be pushed away from the unstable fixed point and in the opposite direction of the input velocity, until it converges to the new stable fixed point (orange/turquoise trajectories in Fig 4b, “below threshold velocity” panels). Thus, for sufficiently small velocities, the bump will be driven to and will persist at the fixed orientation of a stable fixed point, regardless of initial condition and despite being continually driven by a velocity input.

As the velocity input increases, the orientations of both the stable and unstable fixed points will shift further toward the boundary between the stable and unstable regimes. This, in turn, will enable the bump to move closer to this boundary before getting stuck at the stable fixed point. At a particular threshold velocity, the two fixed points will meet at the boundary (Fig 4b, “threshold velocity” panel). Just above this velocity, the bump will be pulled toward the boundary by the stable fixed point, only to be pushed away by the unstable fixed point upon reaching that boundary. This condition enables the bump to smoothly transition between these two regimes without getting stuck. The threshold input velocity *v*_thresh_ for which this occurs depends on how quickly the fixed points approach the boundary between stable and unstable regimes (Fig 4c, left panel), which can be expressed in terms of the net drift speed | *λ*_*d*_|:

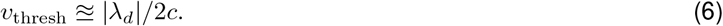

In the limit that the local excitation approaches an optimal value, the net drift speed will go to zero, and the network will be able to integrate infinitesimally small inputs. For intermediate values of local excitation, the net drift speed is moderately high (Fig 4c, right panel), and thus the system will require moderately large input velocities for the bump to move continuously (Fig 4e).

Above this threshold velocity, the orientation of the stable fixed point will shift *ahead* of the stable regime, and the orientation of the unstable fixed point will shift *behind* the unstable regime (Fig 4b, “above threshold velocity” panels). As a result, the bump will be driven by fixed points that it can never reach. In the stable regime, the bump will be pulled from ahead toward the stable fixed point; however, as soon as the bump crosses the boundary into the unstable regime, it will be pushed from behind by the unstable fixed point. This push-and-pull dynamic causes the bump to speed up and slow down as it moves through these two regimes; the closer the fixed points are to this boundary (Fig 4d, left panel), the more pronounced this effect will be (Fig 4d, right panel), and the more nonlinear the resulting bump trajectories will be (Fig 4f).

Because the bump slows down within the stable regime and speeds up within the unstable regime, the slowest and fastest bump velocities *ν*_min_ and *ν*_max_ occur at the boundaries between these regimes, and depend on the relative difference between the threshold and input velocities (see *Methods* |*Model Analytics* |*Locations of Fixed Points in Velocity-Driven Regime* for more details):

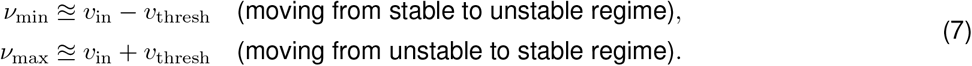

In the limit that the local excitation approaches an optimal value, the threshold velocity *v*_thresh_ will go to zero, and the bump will continuously move at the rate of the input velocity *v*_in_. As the local excitation is tuned away from these optimal values, the threshold velocity will increase, and thus so too will the range of observed bump velocities (Fig 4g). However, because this range depends only on the fixed threshold velocity, its relative effect will be smaller as the input velocity increases.

### Tolerances about optimal parameter values scale linearly with network size

The analytic tractability of this network enables us to characterize performance as we tune the system toward different optimal values of local excitation. As the local excitation approaches an optimal value, (1) the bump will drift more slowly in the absence of input (Eq. 5; Fig 3d, right panel), (2) the network will be able to integrate increasingly small inputs (Eq. 6; Fig 4c, right panel), and (3) the integration of larger inputs will become increasingly more linear (Eq. 7; Fig 4d, right panel). As the size of the network increases, there are increasingly more values of local excitation that achieve optimal performance (Fig 2d), and the degradation of performance away from these values is less severe (Fig 5a-b). This degradation determines how precisely the local excitation should be tuned to meet a threshold level of performance (Fig 5c). For small threshold values, we can analytically determine the widt of the interval about each optimal value of local excitation 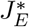 for which the performance is within this threshold; we define this to be the tolerance tol 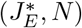 (see *Methods* |*Model Analytics* |*Tolerance in Tuning*):

**Figure 5:**
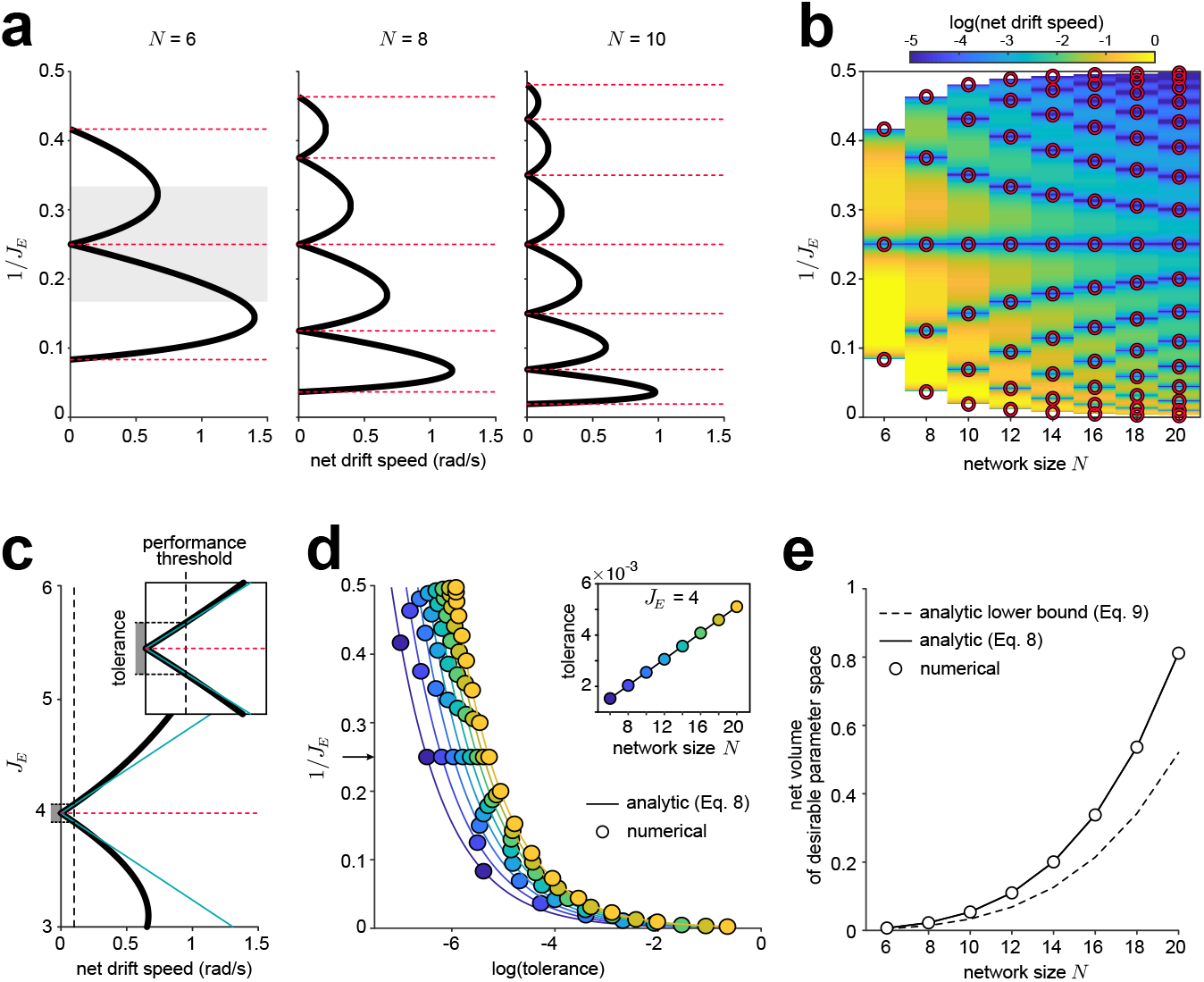
Performance varies as a function of local excitation and network size. **a)** Net drift speed as a function of local excitation for three different network sizes (analogous to the right panel of Fig 3d, shown here for all optimal and intermediate values of local excitation). As network size increases, there is an increasing number of optimal values of local excitation for which the bump does not drift (red dashed lines). Between these values, the network achieves better performance for larger network sizes (larger *N*). Note that the net drift speed appears to increase more rapidly with *J*_*E*_ about larger optimal values of local excitation (smaller values of 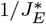); this is purely a product of our vertical axis choice of 1*/J*_*E*_. In fact, we find that the net drift speed increases more slowly with *J*_*E*_ about larger optimal values of local excitation (smaller values of 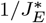); see panel **(d). b)** Same as **(a)**, but for all evenly-sized networks between *N* = 6 and *N* = 20. Red circular markers indicate optimal values of local excitation. The color denotes the log of the net drift speed, with darker blue indicating slower (and hence better) drift rates. **c)** Same as gray highlighted region in left panel in **(a)**, but the net drift speed (black line) is plotted as a function of *J*_*E*_, rather than 1*/J*_*E*_. We use the local change in net drift speed with respect to *J*_*E*_ (turquoise lines) to estimate the tolerance around each optimal value of local excitation (length of gray shaded region) that will achieve performance below some threshold (vertical dashed black line, illustrated for a threshold of 0.1 rad/s). Inset shows a zoomed-in version of the main panel. **d)** For a given network size *N* (indicated by color), larger values of local excitation (smaller values of 1*/J*_*E*_) have higher tolerance than smaller values of local excitation (larger values of 1*/J*_*E*_). Solid lines mark the analytic tolerance given in Eq. 8; filled circles indicate the numerically-estimated tolerance about each optimal value of local excitation. Computed for a threshold value of 0.001 rad/s. Inset shows the scaling with *N* for *J*_*E*_ = 4 (the only optimal value of local excitation that remains unchanged as we increase the size of networks with even numbers of neurons). See *Methods* |*Model Simulations* |*Evaluating Performance with System Size* for more details. **e)** The net volume of parameter space that achieves a desired performance threshold (estimated by summing the tolerance across all optimal values of local excitation for a given network size *N*) increases faster than *N* ^2^. Computed analytically via Eq. 8 by summing over all optimal values of local excitation (solid black line), and estimated numerically by summing over all values shown in panel **(d)**. The analytic lower bound given in Eq. 9 is shown for comparison (gray dashed line).

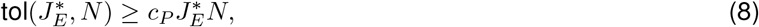

where *c*_*P*_ is a constant that depends on the specific performance measure and desired performance criteria. Larger networks not only produce more optimal values of local excitation, but we find that they also exhibit higher tolerance around each optimal value (Fig 5d; note color progression). This can be seen most clearly for 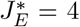 4, which is an optimal value of local excitation for all evenly-sized networks: the tolerance about this value increases linearly with network size (Fig 5d, inset). However, the tolerance also increases with the strength of local excitation; as a result, networks of a fixed size are more robust to variations in tuning about larger versus smaller optimal values of local excitation (Fig 5d; note trend for each color).

When summed across all optimal values of local excitation, Eq. 8 allows us to estimate the net volume of parameter space that can achieve a desired performance threshold:

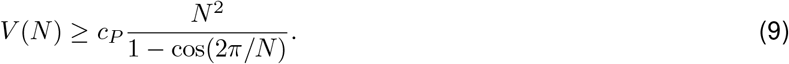

As a consequence of both the number and precise values of optimal excitation, the net volume of desirable parameter space increases at least quadratically with network size (Fig 5e).

Together, these results show that it is possible for a small, discrete attractor network to recover the performance of an infinitely large attractor network when the local excitation in the network is properly tuned. They further show how robustly this performance can be maintained as the local excitation is varied away from these optimal values.

## DISCUSSION

Continuous attractor networks have provided a common theoretical framework for studying a wide range of computations involved in working memory [2, 3, 11, 12], navigation [5, 13–16, 29, 31], and motor control [8, 10, 17]. Across all of these different task domains, this framework has historically relied on an assumption that the underlying networks have large numbers of neurons in order to ensure smooth and accurate dynamics. However, there is growing evidence that similar classes of computations might be performed in much smaller brains that involve far fewer neurons [6, 36, 37, 39, 40, 45–49]. Here, we asked to what extent network size limits the performance of attractor networks [2, 50, 51], and whether small networks can be appropriately tuned in order to overcome these limitations. We focused on a class of attractor networks that maintain a persistent internal representation of a single circular variable like orientation, and that can update this representation by integrating an internal signal like angular velocity. In the limit of infinite numbers of neurons, these ring attractor networks generate a continuous ring manifold along which the population activity will smoothly and accurately evolve. When the number of neurons is finite, we found that networks with as few as 4 neurons could recover this continuous ring attractor manifold, so long as the tuned component of the connectivity (what we refer to as local excitation) is precisely chosen. In the threshold linear networks studied here, this manifold emerges as a set of line attractor manifolds that govern the dynamics of active subsets of neurons in the network, and that are stitched together to generate a complete ring manifold. The resulting population activity can persist at any orientation in the absence of input, and it can smoothly integrate small and large velocity inputs.

Together, these results suggest that very small networks can achieve levels of performance that were thought to require large numbers of neurons. However, this performance comes at the cost of finely tuning the local excitation in the network to one of a discrete number of optimal values. Our biological inspiration for exploring small attractor networks was the fly head direction circuit [6, 37, 39, 40], but this system’s performance during free behavior is unknown, as it has thus far only been probed under head-fixation (see *Methods* |*Experimental Setup* |*Spherical Treadmill System*). Importantly, some inaccuracies in the measured performance of the HD circuit may be attributable to its inputs, that is, to errors in computations of angular velocity and not of its integration. We note that there are many differences between the fly circuit and the simple model we explore here, and some of these differences may provide as-yet-undescribed mechanisms to overcome the potential problems of discreteness in the system. For example, a potential substrate for tuning local excitation may be the synaptic contacts that the fly’s HD neurons make between themselves in different substructures of the CX [20, 37]. Some of these and other fine-scale details of synaptic connectivity in the network have not been incorporated into any existing rate models [36, 40] or spiking neuron models [45–48] of the circuit. In addition, these modeling efforts have focused on capturing the dynamics of the circuit without incorporating the biophysical properties of its neurons, and, in most cases, with only a subset of the excitatory and inhibitory cell types likely involved in generating the dynamics. Although the receptor and transmitter profiles of the relevant neurons are known [37], further experiments are required to assess how much the intrinsic properties of neurons in the fly HD circuit might account for some of the system’s properties, such as persistent activity, as has been reported in the mammalian HD system [52]. Thus, while our work provides a clear indication that small-sized networks can, with appropriate tuning, perform as ring attractor networks, further experiments are needed to investigate their implementation at the level of cellular and synaptic mechanisms in real circuits.

Importantly, large ring attractor networks also suffer from the problem of fine-tuning, where noise in the connectivity —arising, for example, from heterogeneity in synaptic or cellular properties—can yield bumpy energy landscapes similar to those generated here (Fig 2e). Several different mechanisms have been proposed to combat this issue, including homeostatic synaptic scaling [53] and synaptic facilitation [54]. These same mechanisms might also be effective in the small networks studied here, where—in addition to fine-tuning the profile of the connectivity—the overall strength of local excitation must be tuned to one of a discrete number of optimal values. Away from these optimal values, network dynamics are governed by two distinct sets of unstable and stable linear regimes in which the population activity is pushed from or pulled toward discrete fixed points. We identified three properties of regimes within each set that govern network performance: the angular width of each regime, the locations of fixed points within each regime, and the speed at which the bump is pushed from or pulled toward each fixed point. Varying the strength of local excitation in the network alters the balance between the two sets of regimes, such that improving performance in one set of regimes worsens performance in the other. However, as the local excitation approaches an optimal value, the relative impact of the set of worse-performing regimes shrinks to zero, such that overall performance is dominated by the set of better-performing regimes. In the limit that local excitation is precisely tuned to an optimal value, the dynamics are governed entirely by the set of better-performing regimes, which in the same limit become a set of line attractors.

This analysis relied on characterizing the behavior of threshold-linear networks in terms of a separation between different linear dynamical regimes. This separation of linear dynamics has recently been used to infer the underlying connectivity of biological networks [55], and to design different connectivity motifs that use transitions between linear regimes to keep count, to coarsely represent different positions, or to generate distinct dynamical patterns [56, 57]. Here, we showed how the precise tuning of interactions within a single connectivity motif shapes the properties of these linear regimes, and how these properties in turn impact the accuracy of integration in the network. We found that certain regions of parameter space yield more accurate integration, and among these “good” parameter regions, some are more robust than others. Specifically, we found that larger optimal values of local excitation, which result in narrower, more sharply-tuned activity bumps, are more robust to variations in tuning. Previous studies of attractor networks have shown that more sharply-tuned activity bumps are also more robust to noise [2, 51], suggesting that stronger local excitation might jointly enhance tuning and noise robustness.

Our results relied on specific assumptions about network connectivity and dynamics. We assumed local cosine-tuned excitation and broad uniform inhibition, but ring attractor manifolds can be generated with different hand-tuned [21, 23, 28, 32, 58] or learned [59] connectivity structures. To perform velocity integration, we relied on velocity-dependent weights within a single-ring architecture [23]. Velocity integration can instead be performed using a network of two rings that receive differential velocity input [28], or via the addition of two side rings that inherit heading activity from a center ring and project back to the center ring with a velocity-dependent phase shift [22], as has been observed experimentally [39, 40]. Our formulation approximates this second implementation in the limit that the two side rings have fast neural time constants. Finally, our dissection of network dynamics relied on our choice of a threshold-linear response function, which enabled us to decompose the dynamics into distinct linear regimes [43, 44] and analytically dissect failures of integration within these different regimes. This allowed us to study not only how performance varied as a function of network tuning, but also how network size shapes the tuning precision required to achieve a desired level of performance. In the threshold-linear networks considered here, this precision is limited to the tuned component of the connectivity, but in networks with other nonlinearities, both the tuned and untuned components would need to be precisely chosen (SI Fig S3a). In the absence of such precision, small networks can fail to integrate velocity inputs and can drift in the absence of input. While such performance failures are known to arise in small attractor networks with differing connectivity structures and neural response functions [2, 50, 51], it remains an open question as to how these different design features impact the relationship between tuning precision and performance more broadly.

## ACKNOWLEDGEMENTS

We thank James E. Fitzgerald for useful discussions about characterizing threshold-linear dynamics in terms of linear subspaces. This work was funded by the Howard Hughes Medical Institute.

## AUTHOR CONTRIBUTIONS

MN, SR, and AMH conceptualized the problem, with input from VJ. BKH performed all experiments and data processing. VJ performed data analysis, with primary input from BKH, and additional input from MN, SR, and AMH. MN performed the bulk of the analytics, with contributions from SR and AMH. MN, SR, and AMH performed simulations. MN and AMH wrote the paper, with input and editing from all authors.

## COMPETING INTERESTS

We do not have any competing interests.

## METHODS

### Experimental Setup

#### Fly Preparation for Imaging

We expressed the genetically-encoded calcium indictor GCaMP6f [60] in EPG neurons by crossing GCaMP6f flies (w1118;;PBac[20XUAS-IVS-Syn21-op1-GCaMP6f-p10] in VK00005) to the EPG GAL4 driver line VT25957 [61]. Flies (females, age 8 − 11 d, *n* = 27) were prepared for imaging as previously described [6, 62]. Briefly, flies were anesthetized at 4 C, their proboscis immobilized with wax to reduce brain movements, and their head/thorax fixed to a holder with a recording chamber using ultraviolet glue. To gain optical access to the brain, a section of cuticle between the ocelli and antennae was removed, along with the underlying fat and air sacs. Throughout the experiment, the head was submerged in saline containing (in mM): NaCl (103), KCl (3), TES (5), trehalose (8), glucose (10), NaHCO_3_ (26), NaH_2_PO_4_ (1), CaCl_2_ (2.5) and MgCl_2_ (4), with a pH of 7.3 and an osmolarity of 280 mOsm.

#### Two-Photon Calcium Imaging

Calcium imaging was performed with a custom-built two-photon microscope controlled with ScanImage (Vidrio Technologies; [63]). Excitation of GCaMP6f was generated with an infrared (920 nm), femtosecond-pulsed (pulse width ∼ 110 fs) laser (Chameleon Ultra II, Coherent) with 15 mW of power, as measured after the objective (×20 Olympus XLUMPLFLN, 1.0 numerical aperture, 2.0 mm working distance). Fast Z stacks (eight planes with 6 *μ*m spacing and three fly-back frames) were collected at 10 Hz by raster scanning (128 × 128 pixels; ∼ 60 × 60 *μ*m^2^) using an 8 kHz resonant-galvo system and piezo-controlled Z positioning. Focal planes were selected to cover the full extent of EPG processes in the ellipsoid body. Emitted light was directed (primary dichroic: 735; secondary dichroic: 594), filtered (filter A: 680 SP; filter B: 514/44) and detected with a GaAsP photo-multiplier tube (H10770PB-40, Hamamatsu).

#### Spherical Treadmill System

Following dissection, flies were positioned on an air-supported polyurethane foam ball (8 mm diameter, 47 mg) under the two-photon microscope and allowed to walk. Rotations of the ball were tracked at 500 Hz, as described previously [62]. Behavioral data and imaging timestamps were recorded using WaveSurfer (http://wavesurfer.janelia.org/). For each fly, we collected twelve 1-minute trials during which flies walked in darkness. We note that this situation can never perfectly simulate free walking. Slight asymmetries in tethering flies, and differences in the proprioceptive feedback that a fly might receive when walking on a ball heavier than itself while head-fixed and with its brain exposed, are likely to affect the accuracy of angular velocity inputs that the fly compass receives in our setup.

## Data Analysis

All data analysis was performed in MATLAB (MathWorks Inc., Natick, MA). Some analyses relied on functions from the Circular Statistics Toolbox [64].

### Extracting Bump Orientation and Strength

Each Z-stack was reduced to a single frame using a maximum intensity projection. An ellipse was manually drawn around the perimeter of the ellipsoid body and automatically segmented into 16 equal-area, wedge-shaped regions of interest (ROIs). The number of ROIs was chosen to match the number of anatomically defined ellipsoid body wedges [65]. Activity within each ROI was averaged for each frame, producing 16 ROI time series. For each ROI time series, baseline fluorescence (*F*_0_) was defined as the average of the lowest 10% of samples. Δ*F/F* was computed as 100 × (*F* − *F*_0_)/(*F*_0_), where *F* is the instantaneous fluorescence from the raw ROI time series.

We used the population vector average (PVA) as a measure of bump strength and orientation. PVA was computed by taking the circular mean of vectors whose angles were the ROI’s wedge positions and whose length was equal to the ROI’s Δ*F/F*. The magnitude of this mean resultant vector length was normalized to have a maximum possible length of one.

### Characterizing Bump Drift

To determine bump drift, we first identified periods when flies were standing still (defined as zero rotational and translational velocity), disregarding periods shorter than 300 ms. Drift was computed as the circular distance between bump orientations (PVA phase) at the beginning and end of these periods of standing.

### Characterizing Bump Velocity

We computed the instantaneous angular velocity of the fly as the change in its orientation over 400 ms. To compute bump velocity we used the circular distance between the bump orientations over that period. If the PVA magnitude dropped below 0.25 during the epoch, the 400 ms epoch was discarded from further analysis. We computed the maximum (absolute) angular velocity during this period, and created 20 velocity bins within a range spanning the negative and positive values of the maximum velocity. Similarly, bump orientations in the EB were divided into 32 bins, each π/16 rad wide. Bump velocity as a function of the bump’s position and the fly’s rotational velocity were placed in these bins. If there were fewer than 3 values in a bin, the bin was disregarded; otherwise the bump velocities within the bin were averaged and shown in Fig 1j. Finally, to display average bump velocities corresponding to different rotational velocities and bump orientations across flies (bottom panel of Fig 1j), we normalized the bump velocities by a fly-specific and trial-specific gain before computing binned averages. This gain was computed by fitting a linear model to bump and rotational velocities from each trial.

## Model Overview

### Network Equations

We consider an effective single-ring network of *N* neurons (or equivalently, of *N* computational units; see *Supplemental Information* |*Network Equations*). Neurons are ordered according to their preferred heading *θ*_*j*_, which we take to be evenly spaced by Δ*θ* = 2π*/N* radians. Neurons are recurrently connected according to their preferred headings via a symmetric weight matrix 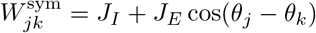, where *J*_*E*_ and *J*_*I*_ parametrize the strength of local excitation and uniform inhibition, respectively (note that *J*_*E*_ and *J*_*I*_ actually correspond to tuned and untuned components of the connectivity; for ease of language, we use local excitation and broad inhibition here and throughout. See *Results* |*Optimally-tuned discrete networks generate a continuum of stable configurations*). Neurons receive velocity input via an asymmetric, velocity modulated weight matrix 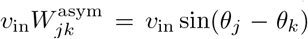; in the main text, we took *v*_in_ *>* 0. Each neuron *j* receives a constant feedforward input *c*_*ff*_ and a net input 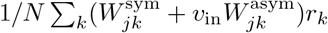 from all other neurons in the network, where the firing rate *r*_*k*_ = (*h*_*k*_) is a nonlinear function of the total input activity *h*_*k*_. For all analyses shown in the main text, we took the nonlinear transfer function *ϕ* (·) to be rectified linear (i.e., *ϕ* (·) = [·]_+_, but see also SI Fig S3 and *Methods* |*Model Simulations* |*Robustness to Changes in the Transfer Function and Recurrent Weights*). The dynamics of each neuron are given by the system of single-neuron equations in Eq. 1; we chose τ = 0.1 s and *c*_*ff*_ = 1.

By applying a discrete Fourier transform to the single-neuron equations, we can express this system of equations in terms of its Fourier modes. After initial transients, only the DC and first-order modes remain, and the resulting dynamical system reduces to set of three equations that govern the dynamics of the orientation *ψ*, amplitude *a* relative to the average input activity, and width *w* of the bump (*Supplemental Information* |*Network Equations* |*Order Equations*); we will refer to these as the system of bump equations.

### Stable Parameter Regime

The system of bump equations will generate a stable bump of activity for certain combinations of *J*_*E*_ and *J*_*I*_ (see *Supplemental Information* |*Fixed Point Solutions* |*Fixed Point Analysis* and SI Fig S1). For all analyses shown in the main text, we first selected a desired value of *J*_*E*_ *>* 2, and we then selected a value of *J*_*I*_ such that it produced a bump of activity whose full amplitude *A* = *H*_0_ + *a* (where *H*_0_ is the average input activity) was at least about *A* = 0.2. To do so, we first uniformly sampled bump orientations *ψ* − [0, 2π) and widths *w ∈* [2π*/N*, 2(*N −* 1)π*/N*), and we used these to calculate the contour *J*_*E*_*f*_even_(*w, ψ*) = 1 using MATLAB’s *contourc*.*m*, where *f*_even_(*w*,) is given by SI Eq. S19 (see also SI Eq. S30 and SI Fig S6). This gave us values 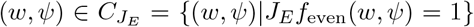 that satisfy the contour equation. We then used these values of *w* and *ψ* to determine an upper bound on *J*_*I*_ given by

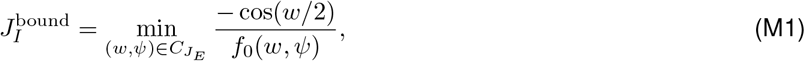

where *f*_0_(*w, ψ*) is given by SI Eq. S18 (see also SI Eq. S32). We then used these same values of *w* and *ψ* to determine a value for *J*_*I*_, given by

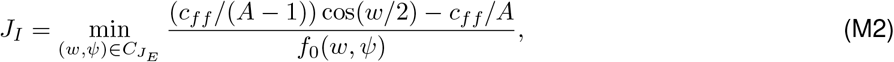

and verified that 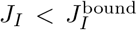. Plugging *A* = 0.2 into Eq. M2 resulted in a bump of activity whose minimum full amplitude was around *A* = 0.2.

## Model Analytics

### Stationary Solutions

To determine the configurations to which the system evolves in the absence of velocity input, we characterized the stationary solutions of the system of bump equations (*Supplemental Information* |*Fixed Point Solutions* |*Fixed Point Analysis*). This allowed us to determine relationships between the bump orientation, relative amplitude, and width that would persistently maintain a stable bump of activity (see SI Fig S6). For a network of *N* neurons that receives no velocity input, the majority of parameter settings will yield two sets of *N* fixed points each—one set will be stable, and the other will be unstable. For a given value of *J*_*E*_, one set will be aligned with the preferred headings {*θ*_*j*_}, and the other set will be aligned precisely between the preferred headings; the second and fourth columns of Fig 2e highlight examples for which the unstable (second column) and stable (fourth column) sets of fixed points are aligned with the preferred headings. The value of *J*_*E*_ and the parity of *N* (whether the network consists of an even or odd number of neurons) together specify which of these two configurations the network will adopt. When *N* is even and 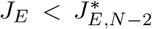 (denoting bumps supported by *N −* 1 and *N−* 2 neurons), the set of fixed points aligned with the preferred headings will be *unstable*. When *N* is odd, the reverse will be true: for 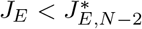, the set of fixed points aligned with the preferred headings will be *stable*. For a given network size *N*, as *J*_*E*_ passes through an optimal value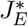, this stability switches. At each of these fixed points, the widths of the stable/unstable bump configurations are determined solely by *J*_*E*_, while their relative amplitudes depend on both *J*_*E*_ and *J*_*I*_.

### Energy Landscape

We derived an energy landscape *E*(*a, w, ψ* ; *J*_*E*_, *J*_*I*_) for the system of bump equations in the absence of velocity input [41, 42] (*Supplemental Information* |*Fixed Point Solutions* |*Energy Landscape*). This function describes the stable configurations to which the system will evolve in the absence of input.

To minimize the curvature of the energy landscape, we first determined the 3 × 3 Hessian matrix of second derivatives of the energy *E* with respect to *a, w*, and *ψ*. When evaluated at the orientations *ψ*^*s*^ of the stable fixed points (see previous subsection), we found that the Hessian reduced to a block diagonal matrix, with a single eigenvector along whose eigenvalue is given by:

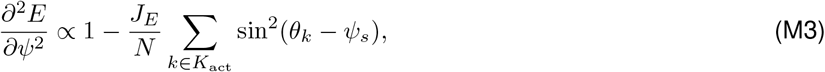

where *K*_act_ denotes the set of indices of the neurons that actively maintain the bump. This eigenvalue quantifies the degree of local curvature as a function of bump orientation *ψ*. For a system of size *N*, there are *N −* 3 values of local excitation *J*_*E*_ for which this eigenvalue goes to zero, and thus for which the energy landscape is locally flat as a function of. These correspond to bump configurations for which the bump is maintained by *N*_act_ *∈* [2,*N −* 2] active neurons:

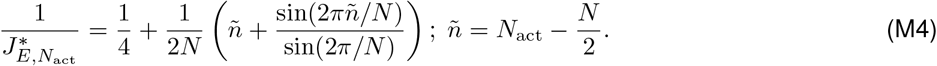

We found that these values of local excitation, which are shown in Fig 2d for a network of size *N* = 6, also ensure that the energy landscape is flat for all bump orientations (as shown in Fig 2e; also see SI Fig S2).

### Leading Eigenvalues of Active Submatrices

In the absence of velocity input, the bump dynamics are governed by the leading eigenvalue of a submatrix of the connectivity (*I* + *W* ^sym^*/N*)*/τ*; this eigenvalue determines the rate at which the bump will drift in the absence of input. When the local excitation *J*_*E*_ is optimally-tuned (i.e., 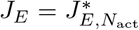), the bump of activity will be maintained by a fixed number of active neurons *N*_act_ *∈* [2, …, *N −* 2]. For each distinct value of *N*_act_, there is thus a distinct *N*_act_ × *N*_act_ submatrix of the connectivity whose single leading eigenvalue determines the drift dynamics. Away from these optimal values of local excitation, the bump of activity will be maintained by either *n* or *n* +1 active neurons (see SI Eq. S47). The drift dynamics are then governed by the leading eigenvalues of the corresponding *n* × *n* and (*n* + 1) × (*n* + 1) active submatrices.

To determine these dynamics, we analytically determined the rates of bump drift in the stable and unstable regimes, which are given in Eq. 3 (see *Supplemental Information* |*Performance of Non-optimal Solutions* |*Dynamics in the Absence of Input Velocity*, and in particular SI Eqs. S51 and S53). We then compared these analytically-derived drift rates to the leading eigenvalues that we computed numerically by directly diagonalizing active submatrices of the connectivity (using the MATLAB function *eig*.*m*); this comparison is shown in SI Fig S4.

### Widths of Stable and Unstable Regimes

In the absence of input, the widths of the stable and unstable regime can be determined analytically by finding the orientation at which the bump transitions from unstable to stable dynamics as it drifts away from an unstable fixed point. This reduces to matching two exponential equations that govern the dynamics of the bump orientation in the two regimes (with drift rates *λ*_*u*_ and *λ*_*s*_, respectively), and that must tend toward the orientations of the unstable and stable fixed points as *t →* − *∞* and *t →* +*∞*, respectively. The resulting widths of each regime are given by Eq. 4 and shown in the middle panel of Fig 3d, and they are centered on the orientations of the stable and unstable fixed points. Given a stable fixed point at *ψ*= *ψ*^*s*^ and an unstable fixed point at *ψ*= *ψ*^*u*^ = *ψ*^*s*^ + π*/N*, the resulting equation for the bump can then be written as (see SI Eqs. S58-S59):

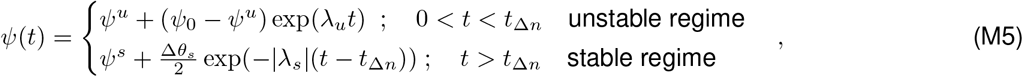

where *ψ*^*s*^ + Δ*θ*_*s*_/2 *< ψ*_0_ *< ψ*^*u*^ is the initial orientation of the bump, and *t*_Δ*n*_ = (1*/λ*_*u*_) log (Δ*θ*_*u*_/(2(*ψ*^*u*^ − *ψ*_0_))) is the time when the bump orientation crosses from the unstable regime into the stable regime. See *Supplemental Information* |*Performance of Non-optimal Solutions* |*Dynamics in the Absence of Input Velocity* for more details.

### Drift in the Absence of Input

To measure the net bump drift, we analytically computed the time *τ*_*d*_ that it takes for the bump to drift from within *ε*_*u*_ of an unstable fixed point to within *ε*_*s*_ of a stable one. We chose *ε*_*u*_ = Δ*θ*_*u*_/2*e* and *ε*_*s*_ = Δ*θ*_*s*_/2*e*, such that the bump covered an angular distance of *Δ ψ*_*d*_ = (1 *−* 1*/e*) Δ*θ/*2 in the time *τ*_*d*_. We then measured the net drift speed as *Δ ψ*_*d*_*/τ*_*d*_ (see SI Eqs. S65-S68).

### Small Velocity Approximation

In the presence of velocity input, the bump dynamics will be governed by the leading eigenvalue of a submatrix of the full connectivity (*−I* + (*W* ^sym^ + *v*_in_*W* ^asym^)*/N*)*/τ*. The asymmetric component of this connectivity is modulated by the input velocity *v*_in_, and introduces a velocity-dependent correction to the eigenvalue *λ*_0_ of the symmetric connectivity (*−I* + *W* ^sym^*/N*)*/τ* (SI Fig S5):

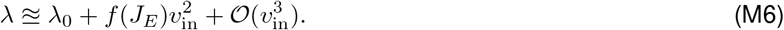

For sufficiently small input velocities, we can approximate the leading eigenvalues *λ*_*u*_ and *λ*_*s*_, and thus the corresponding widths of the unstable and stable regimes, as being equal to their values in the absence of velocity input (see *Methods* |*Model Analytics* |*Leading Eigenvalues of Active Submatrices* and *Methods* |*Model Analytics* |*Widths of Stable and Unstable Regimes*). All analytic results shown in Fig 4b-d,g were generated under this assumption. This approximation breaks down as the input velocity increases, and it breaks down more quickly for smaller values of local excitation (as shown in the right panel of Fig 4d; see also SI Fig S5a).

### Locations of Fixed Points in Velocity-Driven Regime

Although we can approximate the rates and width of the stable and unstable regimes as remaining unchanged for sufficiently small velocity input, we cannot make the same approximation for the orientations of stable and unstable fixed points. We will therefore treat the stable and unstable fixed point orientations as functions of *v*_in_: *ψ*^*s*^ = *ψ*^*s*^(*v*_in_), *ψ*^*u*^ = *ψ*^*u*^(*v*_in_), respectively. The orientation of the stable and unstable fixed points found in the absence of velocity input will then be given by *ψ*^*s*^(0) and *ψ*^*u*^(0), respectively. To determine how the orientations of these fixed points shift with velocity, we repeated the analyses described in *Methods* |*Model Analytics* |*Widths of Stable and Unstable Regimes*, but with a different set of initial conditions (see *Supplemental Information* |*Performance of Non-optimal Solutions* |*Dynamics in the Presence of Small Input Velocity* for details). Given a bump that begins at a stable fixed point at *ψ*= *ψ*^*s*^(0) in the absence of input, and given an initial velocity *v*_in_, the bump will be driven to a new stable fixed point at an orientation *ψ*^*s*^(*v*_in_) = *ψ*^*s*^(0) + *v*_in_/| *λ*_*s*_| as *t → ∞*. In the limit that *t →* − *∞*, the bump will be driven to (and hence, in forward time, away from) an unstable fixed point at an orientation *ψ*^*u*^(*v*_in_) = *ψ*^*u*^(0) *v*_in_/ _*u*_. Over an interval *ψ ∈* [*ψ*^*s*^(0) *θ*_*s*_/2, *ψ*^*u*^(0) + *θ*_*u*_/2], the resulting equation for the bump can be written as (see SI Eqs. S75-S76):

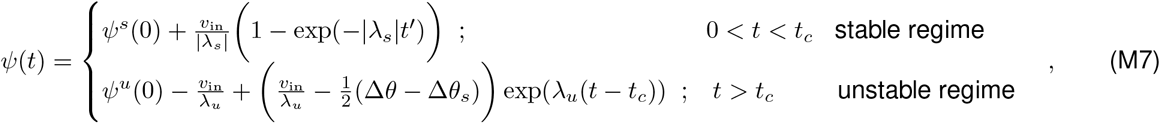

where *t*_*c*_ = (1/| *λ*_*s*_|) log(1/(1 − Δ*θ*_*s*_| *λ*_*s*_|/2*v*_in_)) is the time when the bump orientation crosses from the stable regime into the unstable regime.

At the threshold velocity given in Eq. 6, the two fixed points will meet at the boundary between regimes; this is the minimum velocity needed for the bump to move continuously. Below this velocity, the bump will be driven away from the unstable fixed point in the unstable regime, and toward a stable fixed point in the stable regime. Above this velocity, the stable and unstable fixed points will still drive the bump dynamics, but their orientations will move outside of their respective regimes. The minimum and maximum bump velocities, ν_min_ and ν_max_ (given by Eq. 7), can be computed analytically from Eq. M7 by evaluating the time derivative of (*t*) at the boundary from the stable to the unstable regime, and vice versa. We used these minimum and maximum velocities to define the linearity of integration as ν_min_/ν_max_. See *Supplemental Information* |*Performance of Non-optimal Solutions* |*Dynamics in the Presence of Small Input Velocity* for more details.

### Tolerance in Tuning

To determine how precisely the local excitation must be tuned to achieve a criterion level of performance, we first computed the derivative of each performance measure as a function of local excitation, evaluated at an optimal value; we denote this 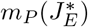 (see SI Eqs. S84-S87). This slope give us a local linear estimate of how quickly the performance degrades away from an optimal value of local excitation. Because each performance measure can be expressed as a function of the net drift speed | *λ*_*d*_|, computing this slope reduced to computing 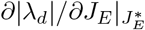. Given a criterion for the system to be within 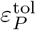 of optimal performance for a performance measure *P*, the tolerance about a given optimal value 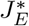 can then be computed as 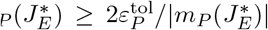 (where ≥ indicates that this is a lower bound on the tolerance, as the linear slope will overestimate the rate of degradation of performance; see SI Eq. S101).

To determine the volume of parameter space that can meet this desired performance, we summed the tolerance across all optimal values of local excitation for a given network size *N* (see SI Eq. S108). We then approximated this sum by its largest value, which reduced to Eq. 9 in the main text. See *Supplemental Information* |*Performance of Non-optimal Solutions* |*Degradation of Performance as a Function of Local Excitation* for more details.

## Model Simulations

### Overview

All simulations that we performed used MATLAB’s ODE solver *ode45*.*m* with an integration timestep of Δ*t* = 0.01 s. We first initialized the network to generate a bump of activity at a given orientation *ψ*. Using this as the initial condition for the network, we then simulated the single-neuron dynamics in Eq. 1, and we performed a discrete Fourier transform using MATLAB’s *fft*.*m* function to extract the bump dynamics as a function of the single-neuron dynamics (see SI Eq. S16). When simulating angular velocity integration, we first determined the velocity scaling that would generate a comparable rate of bump movement for a given (constant) velocity input (see *Methods* |*Model Simulations* |*Velocity-Driven Dynamics*), and then we simulated the network dynamics in response to this scaled input.

### Parameter Choices

All results shown in Fig 2-4 were generated using networks of size *N* = 6. Fig 2e-h was generated using *J*_*E*_ = [12, 6, 4, 3, 2.4] (these values are evenly spaced as a function of 1*/J*_*E*_). For each value of local excitation, we simulated the network dynamics in the presence (Fig 2f, upper row) and absence (Fig 2f, lower row) of velocity input. For the former, we used 10 evenly-spaced velocity values between (and including) 0.2 and 2.0 rad/s, scaled as described below (see *Methods* |*Model Simulations* |*Velocity-Driven Dynamics*).

Figs 3-4 were generated using values of local excitation between two optimal values, 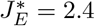 and 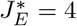. Fig 3b was generated using 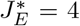. Figs 3c and 4b were generated using *J*_*E*_ = 3. Figs 3f and 4e-g were generated using *J*_*E*_ = [3.6, 3, 2.57] (again, these values are evenly spaced as a function of 1*/J*_*E*_). Comparisons between simulations and analytics in Figs 3d and 4c,d were generated using 17 evenly-spaced values between (but not including) 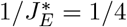 and 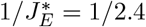 (these values of local excitation include those used in other panels of Figs 3-4).

Fig 4 was generated using a range of different values of the input velocity *v*_in_, scaled as described below. Figs 4a and 4b were generated using velocity values spaced with respect to the threshold velocity *v*_thresh_ = 0.6545 rad/s for a network with *J*_*E*_ = 3: *v*_in_ = [0.209, 0.509, 0.6545, 0.8, 1.1] rad/s. The velocity-dependent orientations of the stable and unstable fixed points in the left panels of Figs 4c and 4d were generated using velocity values in this set: we used *v*_in_ = 0.209 rad/s and *v*_in_ = 0.8 rad/s, respectively. The right panel of Fig 4d was generated using 5 evenly-spaced values between (and including) *v*_in_ = 0.8 rad/s and *v*_in_ = 1.6 rad/s. Finally, Figs 4e and 4f were generated using 10 evenly-spaced values of *v*_in_ between (and including) 0.1 and 1.0 rad/s, and Fig 4g was generated using *v*_in_ = 0.8 rad/s.

### Drift in the Absence of Input

For simulations of bump drift, we simulated the network with the velocity input set to zero. To illustrate drift trajectories for different values of *J*_*E*_ (as shown in Fig 3b,c,f), we initialized the bump at 6 evenly-spaced orientations between (and including) 0 and π*/N*, and we simulated the evolution of the bump for 3 s. To illustrate the stable and unstable portions of these trajectories (as shown in the right panels of Fig 3c,f), we decomposed a single simulated trajectory into portions that lie within the stable and unstable regimes, and we compared these segments to the trajectories that we derived analytically using Eq. M5 (see *Methods* |*Model Analytics* |*Widths of Stable and Unstable Regimes*).

### Measuring Net Drift Speed

To measure the net drift speed (as described in *Methods* |*Model Analytics* |*Drift in the Absence of Input*), we initialized the bump at an orientation *ψ*^*u*^ − *ε*_*u*_ (where *ψ*^*u*^ is the orientation of an unstable fixed point, and *ψ*^*u*^ = π*/N* for the value of *J*_*E*_ used in Fig 3c; see *Methods* |*Model Simulations* |*Parameter Choices*), and we simulated the network dynamics until the bump reached an orientation *ε*_*s*_. We set *ε*_*u*_ = *θ*_*u*_/2*e* and *ε*_*s*_ = *θ*_*s*_/2*e*, where Δ *θ*_*u,s*_ were computed as described in *Methods* |*Model Analytics* |*Widths of Stable and Unstable Regimes*. We used the time it took for the bump to reach this orientation as the measure of the net drift timescale *τ*_*d*_, and we used *Δ ψ*_*d*_*/τ*_*d*_ as a measure of net drift speed, where *Δψ*_*d*_ = (1 *−* 1*/e*) Δ*θ/*2 is the angular distance traveled by the bump in the time *τ*_*d*_. Fig 3d compares the net drift speed from simulations to that obtained analytically for different values of *J*_*E*_.

### Velocity-Driven Dynamics

For simulations of angular velocity integration, we injected a constant velocity input throughout the duration of the simulation. To enable a comparison to analytic predictions, we scaled the input velocity such that the rate of movement of the bump matched the input velocity at an input of *v*_in_ = 50 rad/s. To this end, we determined the best-fitting linear trajectory that minimized the absolute deviation from the bump trajectory over a time window of *t* = 6 s, and we used the slope of this linear trajectory to scale all other input velocities injected into the network. We performed this scaling separately for each set of network parameters (i.e., for each choice of (*J*_*E*_, *J*_*I*_)). All velocity values described in simulations were scaled in this way.

### Measuring Threshold Velocity

To measure the threshold velocity required to move the bump continuously (as shown in the right panel of Fig 4c), we first analytically computed the threshold velocity as described in *Methods* |*Model Analytics* |*Locations of Fixed Points in Velocity-Driven Regime*. We then chose 50 evenly-spaced input velocity values between (and including) *v*_thresh_ 0.05 rad/s and *v*_thresh_ + 0.05 rad/s. We initialized the bump at the orientation of a stable fixed point (here, at *ψ*^*s*^ = 0), and we then simulated the network dynamics in response to each velocity individually. We determined the minimum of these velocities that would move the bump beyond an orientation of π*/N* within a time interval of 10 s. The right panel of Fig 4c compares this simulated value to the value obtained analytically.

### Measuring Linearity of Integration

To measure the linearity of integration from simulations, we simulated the bump trajectory for different constant input velocities (as described above in *Methods* |*Model Simulations* |*Overview*). For each input velocity, we determined the time *t*_*c*_ when the bump orientation *ψ* crossed from the stable into the unstable regime or vice-versa; these times were used to compute the minimum and maximum velocities, respectively (note that we used the analytically-derived boundaries between regimes to determine these crossing times; see *Methods* |*Model Analytics* |*Widths of Stable and Unstable Regimes*). We then determined the bump velocity as ν = (*ψ* (*t*_*c*_ + Δ*t*) − *ψ* (*t*_*c*_ − Δ*t*))/2Δ*t*, where *t* = 0.1 s is the integration timestep used in all simulations. The right panel of Fig 4d compares this simulated value to the value derived analytically (see *Methods* |*Model Analytics* |*Locations of Fixed Points in Velocity-Driven Regime*).

### Evaluating Performance with System Size

To summarize performance as a function of network size (shown in Fig 5a,b), we analytically computed the net drift speed (as described above in *Methods* |*Model Analytics* |*Drift in the Absence of Input*) as a function of local excitation in the range 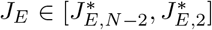 (i.e., between the minimum and maximum optimal values of local excitation, maintained by *N*_act_ = *N* −2 and *N*_act_ = 2 active neurons, respectively). For each optimal value of local excitation, we numerically estimated the tolerance as the range of local excitation values about an optimum for which the net drift speed would be consistently below a fixed performance threshold (we used a threshold value of 0.001 rad/s). We considered only those values of local excitation above the minimum optimal value or below the maximum optimal value to estimate this tolerance; thus, to estimate the tolerance about the minimum and maximum optimal values, we measured the tolerance in only one direction (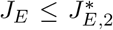 or 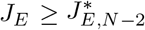), and we doubled this value to use as our estimate. We then compared these tolerance estimates to the analytic lower bound given in Eq. 8, as shown in Fig 5d (also see SI Eq. S101-S107). Finally, we summed these tolerance values (computed numerically or analytically) for each network size *N* to estimate the net volume of parameter space that meets this threshold level of performance, as shown in Fig 5e.

### Robustness to Changes in the Transfer Function and Recurrent Weights

We examined the robustness of the continuous attractor regime to changes in the number of Fourier modes of the recurrent connections in *W* ^sym^, the neuron input-output relationship *ϕ*, and an increase in the dimensionality of the attractor. To this aim, we numerically solved the dynamics of Eq. 1 with *v*_in_ = 0 in two different scenarios. First, we used (i) a Von Mises connectivity profile with concentration parameter *κ* for the recurrent weights 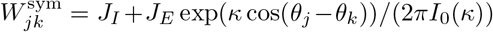, where *I*_0_(*κ*) is the modified Bessel function of order 0; (ii) a smooth nonlinear transfer function *ϕ* (*x*) = log(1 + *e*^*x*^). We numerically solved the dynamics of a network with *N* = 8 units and *J*_*I*_ = 30, with cosine-shaped initial conditions centered at 50 uniformly spaced orientations on the ring (SI Fig S3a). We evaluated the dispersion (circular variance) between the initial and final orientations on the ring for different values of *J*_*E*_, after numerically solving the dynamics for a total time of 500*τ*, where *τ* is the single-neuron time constant. We observed the presence of optimal values of *J*_*E*_ (SI Fig S3a; red), where the network behaved like a continuous attractor, as opposed to other values of *J*_*E*_ (SI Fig S3a; purple, blue) where the discreteness of the solution was evident. The specific values of optimal excitation depend on both the value of *J*_*I*_ (SI Fig S3a; empty circles), and on the strength of constant feedforward input *c*_*ff*_. We next examined the dynamics in Eq. 1 with a recurrent weight profile storing a 2-dimensional toroidal attractor with *N* = 16 neurons, 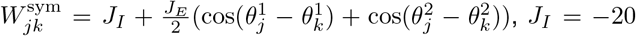, *J*_*I*_ = *−* 20, where the preferred orientations 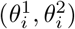 of the units were uniformly spaced on the torus (SI Fig S3b). We similarly observed the presence of an optimal value of *J*_*E*_ for which the dispersion between sub-threshold bumps initialized at 100 different orientations on the torus and the final orientations were close to 0.

## Notes

### Competing Interest Statement

The authors have declared no competing interest.

