## Supplementary Information for "Accurate angular integration with only a handful of neurons"

### SUPPLEMENTARY FIGURES

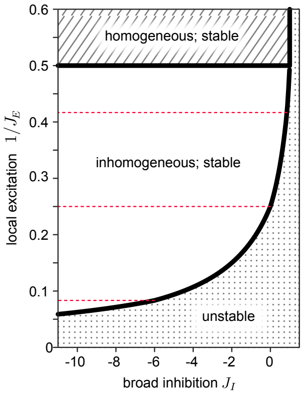


**Figure S1: Parameter regimes that govern the stability of network dynamics.** The stability of the network dynamics depends on both the local excitation $J_E$ and the broad inhibition $J_I$. In the "unstable" regime, the population activity will diverge over time. In the "homogeneous" regime, the network will generate a stable activity profile that is uniform across the entire network. In the "inhomogeneous" regime, the network will generate a stable bump of activity that can persist at a discrete set of orientations in the absence of input. Dashed lines indicate optimal values of local excitation for which the network will generate a set of marginally stable solutions that can persist at any orientation in the absence of input (shown for a network of $N = 6$ neurons).

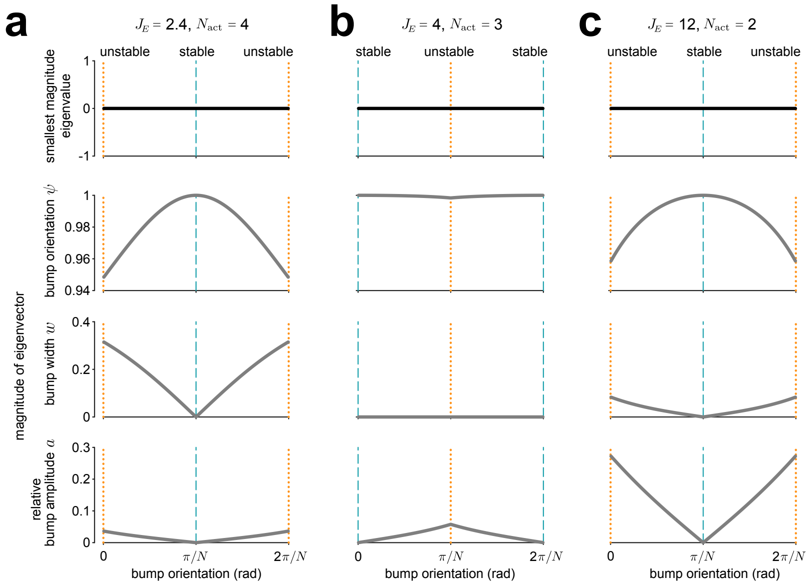


**Figure S2: Flat directions in the energy landscape.** Smallest magnitude eigenvalues (top row) and corresponding eigenvector components (lower three rows) for the Hessian matrix of the energy, computed for all three optimal values of local excitation in a network of size $N = 6$: **(a)** $J_E^* = 2.4$; **(b)** $J_E^* = 4$; **(c)** $J_E^* = 12$. For each optimal value of local excitation, the Hessian has a single zero eigenvalue, indicating the existence of a zero-curvature direction within the energy landscape. The corresponding eigenvectors are purely aligned along $\psi$ (second row) at the orientations of the stable fixed points (teal dashed lines). Away from these orientations, the corresponding eigenvectors involve contributions from $w$ and $a$ (third and fourth rows, respectively).

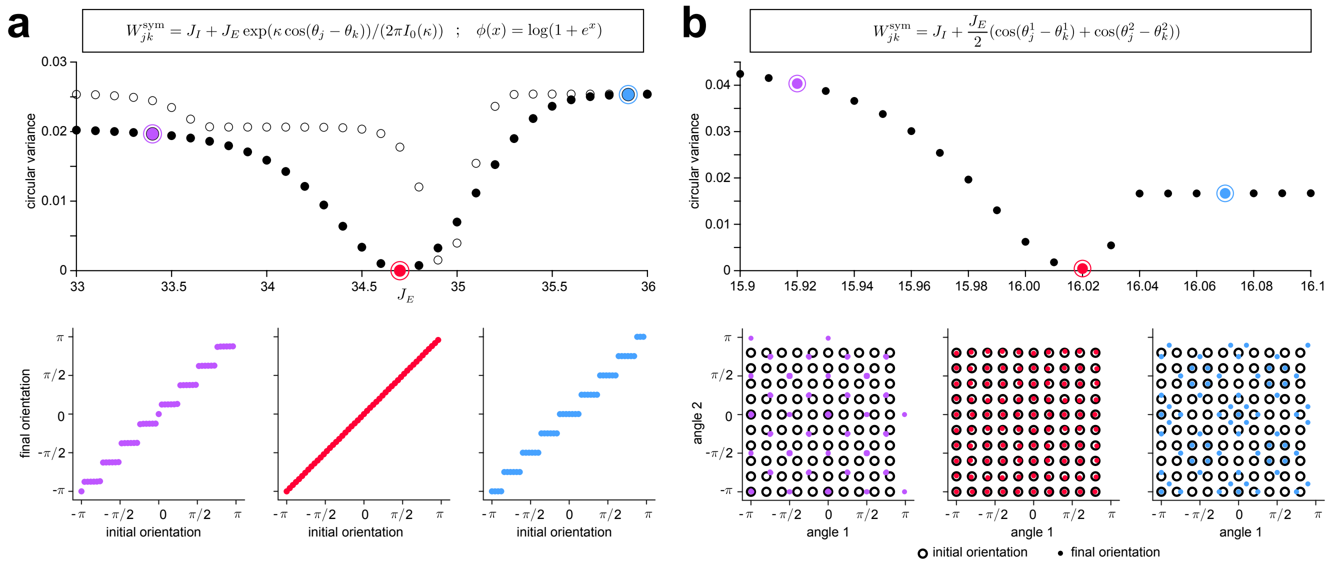


**Figure S3: Robustness to changes in the single neuron transfer function and recurrent synaptic weights.** Comparison between initial and final bump orientations as a function of $J_E$ for **a)** a network of $N = 8$ neurons with a Von Mises weight profile and a smooth nonlinear transfer function, and **b)** a network of $N = 16$ neurons with a recurrent weight profile storing a 2-dimensional toroidal attractor. In both cases, there is an optimal value of $J_E$ for which the circular variance between the initial and final orientations is close to zero (upper panel, red markers), and the bump does not drift (lower center panel). Away from these values of $J_E$, the circular variance increases (upper panel, purple/blue markers), and the bump drifts from its original orientation (lower left/right panels). Filled versus open circles in upper panel of **(a)** correspond to $J_I = -30$ and $J_I = -20$, respectively. See *Methods* |*Model Simulations* |*Robustness to Changes in the Transfer Function and Recurrent Weights* for simulation details.

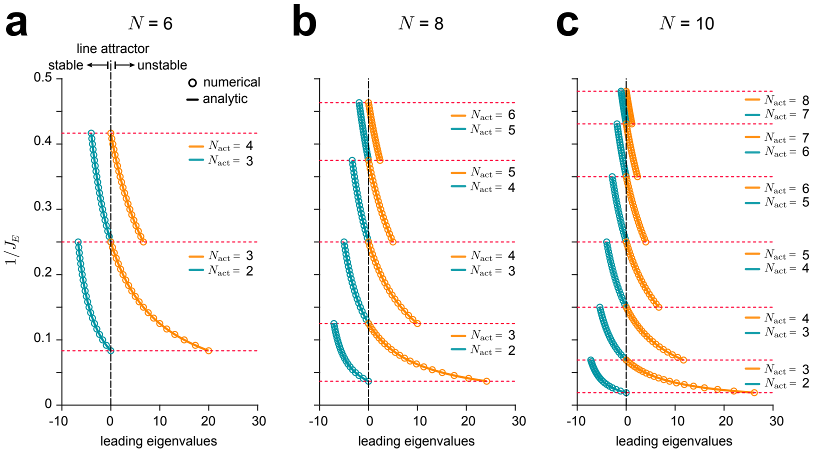


**Figure S4: Leading eigenvalues of active submatrices.** Comparison of analytically- versus numerically- derived eigenvalues (solid lines versus markers, respectively), computed from the active submatrices of the full connectivity $W = (W^{\rm sym}/N - I)/\tau$ in the absence of velocity input. Shown for network sizes **(a)** $N = 6$, **(b)** $N = 8$, and **(c)** $N = 10$. Red dotted lines mark optimal values of local excitation for each network size.

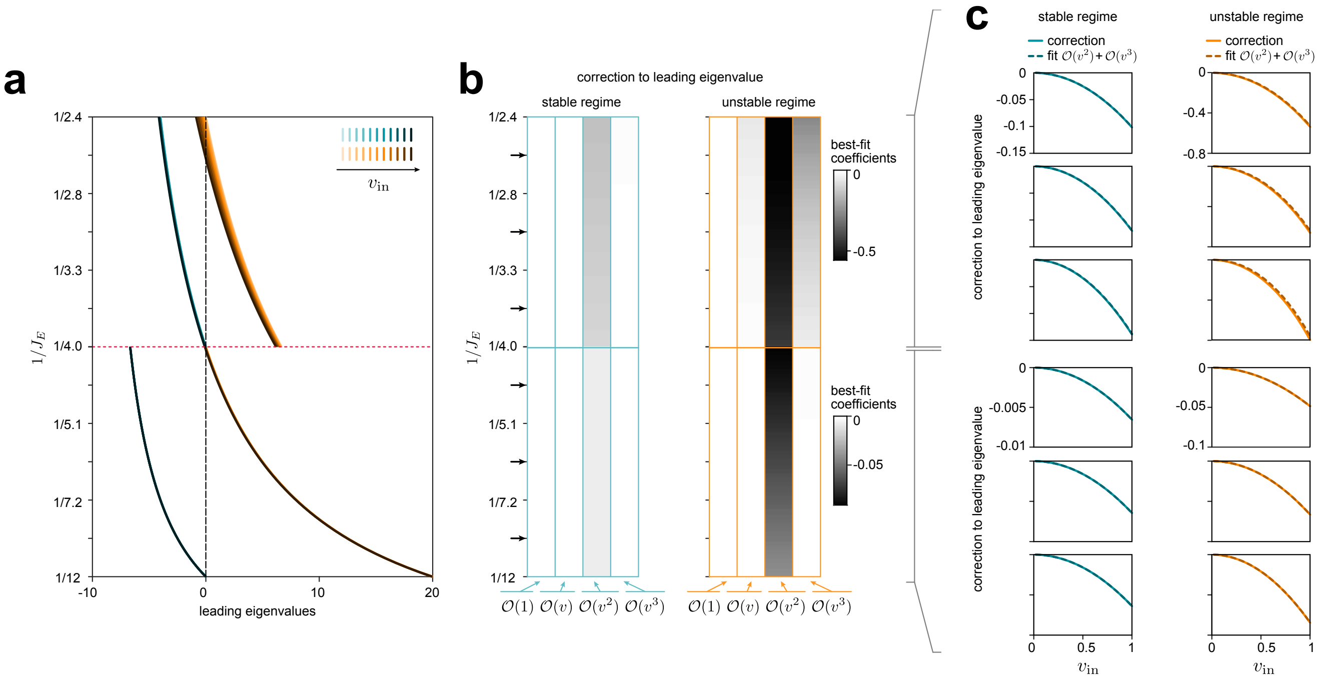


**Figure S5: Velocity correction to leading eigenvalues of active submatrices. a)** Leading eigenvalues $\lambda$ of active submatrices as a function of input velocity $v_{\text{in}}$ for a network of size $N = 6$. Shown for $11$ velocity values evenly spaced between (and including) $v_{\text{in}} = 0$ and $v_{\text{in}} = 1$ rad/s (darker colors indicate higher velocities). Eigenvalues were obtained by numerically diagonalizing active submatrices of the full connectivity $W = ((W^{\text{sym}} + v_{\text{in}} W^{\text{asym}})/N - I)/\tau$. Red dashed line marks an optimal value of local excitation. **b)** Coefficients of the best-fitting $3$rd order polynomial of the velocity correction $\lambda - \lambda_0$ versus input velocity $v_{\text{in}}$, where $\lambda_0$ is the leading eigenvalue of the full connectivity in the absence of velocity input. **c)** Comparison of the velocity correction $\lambda - \lambda_0$ (solid lines) and the best-fitting polynomial (dashed lines), including terms of order $\mathcal{O}(v^2)$ and $\mathcal{O}(v^3)$. Shown for $6$ different values of local excitation marked by arrows in panel **(b)**.

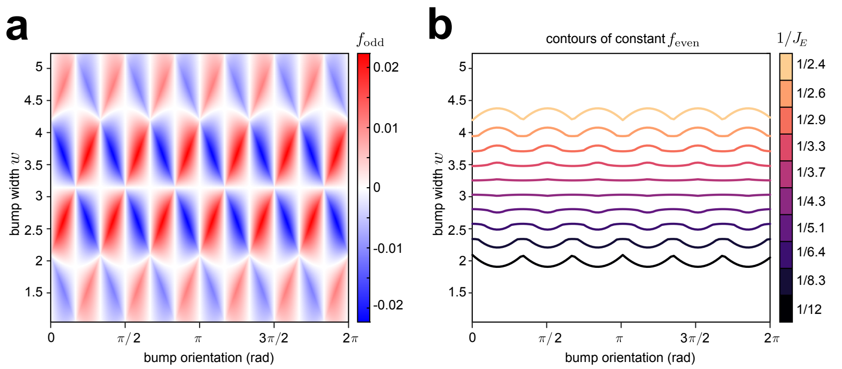


**Figure S6: Visualization of functions $f_{\text{odd}}$ and $f_{\text{even}}$ that govern bump dynamics. a)** Heatmap of the function $f_{\text{odd}}(w, \psi)$ for densely sampled bump widths $w \in [2\pi/N, 2(N-1)\pi/N)$ and orientations $\psi \in [0, 2\pi)$. Red regions correspond to $f_{\text{odd}} > 0$ (which would drive the bump orientation to the right), and blue regions to $f_{\text{odd}} < 0$ (which would drive the bump orientation to the left). White regions indicate $f_{\text{odd}} = 0$, which correspond to potential fixed points $\chi^* = (a^*, w^*, \psi^*)$ at which the bump can stably persist. Note that at $\psi = 1/2(\theta_c + \theta_d)$, $d = c, c+1$, there are white vertical lines that extend across different bump widths, indicating that these values of $\psi$ always result in $f_{\text{odd}}(w, \psi) = 0$, regardless of the value of $w$. See *Supplemental Information* | *Fixed Point Solutions* | *Fixed Point Analysis* for more details. **b)** Contours of constant $f_{\text{even}}(w, \psi)$, shown for $10$ evenly spaced values of $1/J_E$ between (and including) $1/12$ and $1/2.4$. These contours indicate a necessary (but not sufficient) relationship between the bump width $w$ and orientation $\psi$ for stationary bump solutions. Note that this relationship is strongly dependent on $J_E$. See *Supplemental Information* | *Fixed Point Solutions* | *Fixed Point Analysis* for more details.

#### SUPPLEMENTARY INFORMATION - THEORY AND ANALYTICS

#### Network Equations

We consider a simplified system with a triple ring structure, with each ring composed of the same number of computational units, $N$. Here, computational units can be single neurons, or they can by groups of neurons with the same tuning and connectivity. In what follows, we will use "neuron" and "computational unit" interchangeably. We assume that the preferred orientation space $\Theta = \{\theta_1, \ldots, \theta_N\}$ uniformly partitions the full orientation space $[0, 2\pi)$. I.e., $\theta_j = (j-1)\Delta\theta$, $j = 1, \ldots, N$, where $\Delta\theta = 2\pi/N$. Let $h_j$, $j = 1, \ldots, N$, denote the total input activity (e.g. current) to neuron $j$ in the center ring, with preferred orientation $\theta_j$. The input-output relationship for neurons in the center ring is taken to be threshold linear, such that the activity of neuron $j$ is given by $r_j = [h_j]_+$. The center ring system is given by

$$\tau \dot{h}_j = -h_j + c_{ff} + \frac{1}{N}\sum_{k=1}^{N} W^{\text{sym}}_{jk} r_k + h_j^{CW} + h_j^{CCW}, \quad j = 1, \ldots, N, \tag{S1}$$

where dot notation is used for the time derivative, $\tau > 0$ is the neural time constant, $c_{ff} > 0$ is a constant feedforward input to all units in the center ring, and $h_j^*$, $* \in \{CW, CCW\}$, indicates the input received from the respective side rings. The weight matrix $W^{\text{sym}}$ gives the weights of the recurrent connections in the center ring. We take these weights to be given by

$$W^{\text{sym}}_{jk} = J_I + J_E \cos(\theta_j - \theta_k), \quad j, k = 1, \ldots, N. \tag{S2}$$

The parameters $(J_I, J_E)$ can be set such that this system will generate a population profile that qualitatively looks like a discretely sampled "bump" of activity. We will assume the parameters $(J_I, J_E)$ are within the subset $\Omega = \Omega_{J_I} \times \Omega_{J_E}$ of parameter space for which this is the case (shown as the subset marked "inhomogeneous; stable" in SI Fig S1). Note that $\Omega \subset (-\infty, 1) \times (2, \infty)$.

We take the activity in the side ring neurons to be a velocity modulated bump inherited from the center ring that is shifted in orientation space according to whether the ring responds to clockwise (CW) or counter-clockwise (CCW) movements:

$$\tau_v \dot{h}_j^{CW} = -h_j^{CW} + \frac{v^{CW}}{N}\sum_{k=1}^{N} W^{CW}_{jk} r_k, \qquad W^{CW}_{jk} = \cos(\theta_j - \theta_k - \delta), \quad j, k = 1, \ldots, N, \tag{S3}$$

$$\tau_v \dot{h}_j^{CCW} = -h_j^{CCW} + \frac{v^{CCW}}{N}\sum_{k=1}^{N} W^{CCW}_{jk} r_k, \qquad W^{CCW}_{jk} = \cos(\theta_j - \theta_k + \delta), \quad j, k = 1, \ldots, N, \tag{S4}$$

where $\tau_v$ is the neural time constant for the side rings, $v^{CCW} = [v_{\text{in}}]_+$, $v^{CW} = [-v_{\text{in}}]_+$ for angular input velocity $v_{\text{in}}$, and $\delta \in [0, \pi]$ indicates the shift in the inherited bump. Throughout, we will take $\delta = \pi/2$ so that $W^{CW} = -W^{CCW} = W^{\text{asym}}$, where

$$W^{\text{asym}}_{jk} = \sin(\theta_j - \theta_k). \tag{S5}$$

For ease of analytics, we take $\tau_v \to 0$, which reduces the system to a single ring with equations given by

$$\tau \dot{h}_j = -h_j + c_{ff} + \frac{1}{N}\sum_{k=1}^{N} (W^{\text{sym}}_{jk} + v_{\text{in}} W^{\text{asym}}_{jk}) r_k, \quad j = 1, \ldots, N. \tag{S6}$$

**Order Equations.** We will use superscript $B$ ("bump") to distinguish variables that describe the activity of the full population of neurons. Taking the discrete Fourier transform, $H_m^B = \frac{1}{N}\sum_{j=1}^{N} h_j e^{-im\theta_j}$, of (S6) gives us the order equations:

$$\tau \dot{H}_0^B = -H_0^B + \frac{J_I}{N}\sum_{k=1}^{N} r_k + c_{ff}, \tag{S7}$$

$$\tau \dot{H}_1^B = -H_1^B + \frac{1}{2N}\sum_{k=1}^{N} [J_E + v_{\text{in}} i] r_k e^{-i\theta_k}, \tag{S8}$$

$$\tau \dot{H}_m^B = -H_m^B, \quad m = 2, \ldots, \lfloor N/2 \rfloor. \tag{S9}$$

Hence, only the DC mode $H_0^B$ and the first mode $H_1^B$ remain after some initial transients. Note that $\{h_1, \ldots, h_N\}$ is real, and therefore $H_{N-k}^B = \overline{H}_k^B$, where $\overline{z}$ denotes the complex conjugate of $z$. Since $H_1^B \in \mathbb{C}$, we let $H_1^B = \rho^B e^{-i\psi^B}$, and the equations become

$$\tau \dot{H}_0^B = -H_0^B + \frac{J_I}{N}\sum_{k=1}^{N} r_k + c_{ff}, \tag{S10}$$

$$\tau \dot{\rho}^B = -\rho^B + \frac{1}{2N}\sum_{k=1}^{N} [J_E \cos(\theta_k - \psi^B) + v_{\text{in}} \sin(\theta_k - \psi^B)] r_k, \tag{S11}$$

$$\tau \rho^B \dot{\psi}^B = \frac{1}{2N}\sum_{k=1}^{N} [J_E \sin(\theta_k - \psi^B) - v_{\text{in}} \cos(\theta_k - \psi^B))] r_k, \tag{S12}$$

where we assume that we are past any initial transients so that we may ignore (S9). This assumption also allows us to write

$$h_j = H_0^B + H_1^B e^{i\theta_j} + H_{N-1}^B e^{-i\theta_j} = H_0^B + 2\rho^B \cos(\theta_j - \psi^B). \tag{S13}$$

We assume that the parameters $(J_I, J_E) \in \Omega$ so that the population activity appears as a single bump of activity with a proper nonempty subset of neurons active at a single time. Thus, we must have $\rho^B > 0$. We define the amplitude of the bump relative to the average input activity $H_0^B$ to be

$$a^B = 2\rho^B, \tag{S14}$$

and we define the width of the bump $w^B$ by

$$\cos\left(\frac{w^B}{2}\right) = -\frac{H_0^B}{a^B}. \tag{S15}$$

From this definition of bump width, it follows that $h_j > 0$ if and only if $\theta_j \in (\psi^B - w^B/2, \psi^B + w^B/2)$. With these definitions, we can rewrite (S13) as

$$h_j = a^B(\cos(\theta_j - \psi^B) - \cos(w^B/2)). \tag{S16}$$

We can now uniquely describe the population activity by the relative amplitude $a^B$, width $w^B$, and orientation $\psi^B$ of the bump. In what follows, we will refer to this description as the configuration of the population bump, denoted by $\chi^B = (a^B, w^B, \psi^B)$. Note that we can always recover the individual neuron input activities from the bump configuration through (S16). Let $K_{\text{act}}$ denote the set of indices of active neurons in the network. That is,

$$K_{\text{act}} = \{j \,|\, h_j > 0\} = \{j \,|\, \theta_j \in (\psi^B - w^B/2, \psi^B + w^B/2)\}. \tag{S17}$$

We then define the following functions:

$$f_0(w,\psi) = \frac{1}{N}\sum_{k\in K_{\text{act}}}(\cos(\theta_k-\psi)-\cos(w/2)), \tag{S18}$$

$$f_{\text{even}}(w,\psi) = \frac{1}{N}\sum_{k\in K_{\text{act}}}(\cos(\theta_k-\psi)-\cos(w/2))\cos(\theta_k-\psi), \tag{S19}$$

$$f_{\text{odd}}(w,\psi) = \frac{1}{N}\sum_{k\in K_{\text{act}}}(\cos(\theta_k-\psi)-\cos(w/2))\sin(\theta_k-\psi). \tag{S20}$$

We can then rewrite (S10) - (S12) as

$$\tau\dot{a}^B = (-1+J_E f_{\text{even}}(w^B,\psi^B)+v_{\text{in}}f_{\text{odd}}(w^B,\psi^B))a^B, \tag{S21}$$

$$\tau\dot{w}^B = \frac{2}{\sin(w^B/2)}\left(J_I f_0(w^B,\psi^B)+\frac{c_{ff}}{a^B}-(J_E f_{\text{even}}(w^B,\psi^B)+v_{\text{in}}f_{\text{odd}}(w^B,\psi^B))\cos(w^B/2)\right), \tag{S22}$$

$$\tau\dot{\psi}^B = J_E f_{\text{odd}}(w^B,\psi^B)-v_{\text{in}}f_{\text{even}}(w^B,\psi^B). \tag{S23}$$

Note that in (S22), $\sin(w^B/2)\neq 0$ since $w^B\in(0,2\pi)$ by assumption.

#### Fixed Point Solutions

In this section, we will characterize the stationary fixed point solutions that emerge in the absence of input (i.e., for $v_{\text{in}}=0$). In this case, (S6) becomes

$$\tau\dot{h}_j = -h_j+c_{ff}+\frac{1}{N}\sum_{k=1}^{N}W_{jk}^{\text{sym}}r_k, \quad j=1,\ldots,N, \tag{S24}$$

and the order equations, (S21)-(S23), become

$$\tau\dot{a}^B = (-1+J_E f_{\text{even}}(w^B,\psi^B))a^B, \tag{S25}$$

$$\tau\dot{w}^B = \frac{2}{\sin(w^B/2)}\left(J_I f_0(w^B,\psi^B)+\frac{c_{ff}}{a^B}-J_E f_{\text{even}}(w^B,\psi^B)\cos(w^B/2)\right), \tag{S26}$$

$$\tau\dot{\psi}^B = J_E f_{\text{odd}}(w^B,\psi^B). \tag{S27}$$

**Fixed Point Analysis.** We perform a fixed point analysis, setting (S25)-(S27) to zero and solving for $\chi^* = (a^*,w^*,\psi^*)$. We will denote fixed point solutions using a superscript $*$ to distinguish them from more general bump configurations. We will replaced this superscript with $s$ or $u$ once we determine the stability of the fixed points.

Considering first (S27), this gives us

$$f_{\text{odd}}(w^*,\psi^*) = 0, \tag{S28}$$

since $J_E\in\Omega_{J_E}\subset(2,\infty)$ by assumption. Note that whenever $\psi$ is at a preferred orientation $\theta_j$ or is precisely between two preferred orientations, $f_{\text{odd}}(w,\psi)$ will be the sum of an anti-symmetric function about $\psi$, and thus

$$\psi_{cd}^* = \frac{1}{2}(\theta_d+\theta_c), \quad c=1,\ldots,N, \quad d=c,c+1, \tag{S29}$$

will always satisfy (S28), regardless of the value of $w^*$, where here we take the extension of indices given by $c\equiv c \pmod N$. We then expect the system to have at least $2N$ fixed points: $N$ for orientations $\psi_{cc}^*$ at each of the $N$ preferred orientations, and $N$ for orientations $\psi_{c(c+1)}^*$ precisely between consecutive preferred orientations. This assumes, of course, that there exist corresponding values of $a_{cd}^*, w_{cd}^*$ for which (S25) and (S26) will also equal $0$ (which does turn out to be the case). Note that while these $2N$ values of $\psi_{cd}^*$ characterize a set of solutions to (S28), they do not necessarily characterize *all* of the solutions (see SI Fig S6a).

Setting (S25) to zero gives us

$$J_E f_{\text{even}}(w^*,\psi^*) = 1, \tag{S30}$$

since $a^* > 0$ by assumption. Implicit solutions $(w^*, \psi^*)$ to (S30) exist for any $\psi^*$ (given our assumption that $J_E \in \Omega_{J_E}$), indicating a necessary relationship between $w^*$ and $\psi^*$ that is implicitly a function of $1/J_E$ for fixed point solutions (see SI Fig S6b). Specifically, for each $\psi^*_{cd}$ given by (S29), there exists a $w^*_{cd}$ such that (S30) is satisfied. We then need only determine $a^*$ to fully characterize the fixed point solutions to the system. We obtain $a^*$ by setting (S26) to zero and using (S30):

$$a^* = \frac{-c_{ff}}{\cos(w^*/2) + J_I f_0(w^*, \psi^*)}. \tag{S31}$$

Equations (S28),(S30)-(S31) allow us to fully (implicitly) characterize the configuration $\chi^* = (a^*, w^*, \psi^*)$ of the fixed point solutions. Note that, since $a^* > 0$, we must have

$$J_I < \frac{-\cos(w^*/2)}{f_0(w^*, \psi^*)}. \tag{S32}$$

Recall that $f_0(w, \psi)$ is a nonempty sum of positive terms, and thus $f_0(w^*, \psi^*) > 0$.

**Energy Landscape.** The energy of the stationary system is given by

$$E = \sum_{j=1}^{N} \int_0^{h_j} h\phi'(h)\, dh - \frac{1}{2N} \sum_{j,k=1}^{N} W^{\text{sym}}_{jk} \phi(h_j)\phi(h_k) - c_{ff} \sum_{j=1}^{N} \phi(h_j), \tag{S33}$$

where $r_k = \phi(h_k)$ is the input-output function [41, 42], here taken to be threshold linear. Note that $h_j$, $j = 1, \ldots, N$, is bounded, since $(J_I, J_E) \in \Omega$, and thus $E$ is bounded. Since $W^{\text{sym}}$ is symmetric, we have

$$\begin{aligned}
\dot{E} &= \sum_{j=1}^{N} h_j \phi'(h_j)\dot{h_j} - \frac{1}{N} \sum_{j,k=1}^{N} W^{\text{sym}}_{jk} \phi(h_k)\phi'(h_j)\dot{h_j} - c_{ff} \sum_{j=1}^{N} \phi'(h_j)\dot{h_j} \\
&= \sum_{j=1}^{N} \phi'(h_j)\dot{h_j} \left( h_j - \frac{1}{N} \sum_{k=1}^{N} W^{\text{sym}}_{jk} \phi(h_k) - c_{ff} \right) \\
\Rightarrow \dot{E} &= -\tau \sum_{j=1}^{N} \phi'(h_j)(\dot{h_j})^2 \leq 0.
\end{aligned} \tag{S34}$$

Hence, $E$ goes to a minimum wherever $\dot{E} = 0$ as $t \to \infty$. These minima correspond to the stable fixed points of the system. From (S16), we can consider the energy $E$ to be a function of the configuration of the bump. That is, $E = E(\chi)$. Using this, we compute the Hessian of the energy in the space of possible bump configurations (where we drop the superscript $B$ in the partial derivatives for ease of notation):

$$H(E) = \begin{bmatrix} \frac{\partial^2 E}{\partial a^2} & \frac{\partial^2 E}{\partial a \partial w} & \frac{\partial E}{\partial a \partial \psi} \\ \frac{\partial^2 E}{\partial a \partial w} & \frac{\partial^2 E}{\partial w^2} & \frac{\partial^2 E}{\partial w \partial \psi} \\ \frac{\partial^2 E}{\partial a \partial \psi} & \frac{\partial^2 E}{\partial w \partial \psi} & \frac{\partial^2 E}{\partial \psi^2} \end{bmatrix}. \tag{S35}$$

For any $x, y \in \{a, w, \psi\}$, we have

$$\frac{\partial^2 E}{\partial x \partial y} = \sum_{j \in K_{\text{act}}} \left( \frac{\partial h_j}{\partial x} - \frac{1}{N} \sum_{k \in K_{\text{act}}} W^{\text{sym}}_{jk} \frac{\partial h_k}{\partial x} \right) \frac{\partial h_j}{\partial y} + \sum_{j \in K_{\text{act}}} \left( h_j - c_{ff} - \frac{1}{N} \sum_{k \in K_{\text{act}}} W^{\text{sym}}_{jk} h_k \right) \frac{\partial^2 h_j}{\partial x \partial y}. \tag{S36}$$

From (S16), for $j = 1, \ldots, N$, we have

$$\frac{\partial h_j}{\partial a} = \cos(\theta_j - \psi) - \cos(w/2), \qquad \frac{\partial h_j}{\partial w} = \frac{a}{2}\sin(w/2), \qquad \frac{\partial h_j}{\partial \psi} = a\sin(\theta_j - \psi), \tag{S37}$$

$$\frac{\partial^2 h_j}{\partial a^2} = 0, \qquad \frac{\partial^2 h_j}{\partial w^2} = \frac{a}{4}\cos(w/2), \qquad \frac{\partial^2 h_j}{\partial \psi^2} = -a\cos(\theta_j - \psi), \tag{S38}$$

$$\frac{\partial^2 h_j}{\partial a \partial \psi} = \sin(\theta_j - \psi), \qquad \frac{\partial^2 h_j}{\partial a \partial w} = \frac{1}{2}\sin(w/2), \qquad \frac{\partial^2 h_j}{\partial w \partial \psi} = 0. \tag{S39}$$

Evaluating (S36) for different pairs of $x, y \in \{a, w, \psi\}$, we see that both $\frac{\partial^2 E}{\partial a \partial \psi}$ and $\frac{\partial^2 E}{\partial w \partial \psi}$ are summations over an anti-symmetric function about $\psi = \psi^*_{cd}$, for $\psi^*_{cd}$ given in (S29). As a result, if we evaluate the Hessian of the energy at a bump configuration $\chi^B = (a^B, w^B, \psi^*_{cd})$ with orientation $\psi^*_{cd}$, we get

$$H(E)|_{\chi^B} = \left.\begin{bmatrix} \frac{\partial^2 E}{\partial \rho^2} & \frac{\partial^2 E}{\partial \rho \partial \theta_c} & \frac{\partial E}{\partial \rho \partial \psi} \\ \frac{\partial^2 E}{\partial \rho \partial \theta_c} & \frac{\partial^2 E}{\partial \theta_c^2} & \frac{\partial^2 E}{\partial \theta_c \partial \psi} \\ \frac{\partial^2 E}{\partial \rho \partial \psi} & \frac{\partial^2 E}{\partial \theta_c \partial \psi} & \frac{\partial^2 E}{\partial \psi^2} \end{bmatrix}\right|_{\chi^B} = \left.\begin{bmatrix} \frac{\partial^2 E}{\partial \rho^2} & \frac{\partial^2 E}{\partial \rho \partial \theta_c} & 0 \\ \frac{\partial^2 E}{\partial \rho \partial \theta_c} & \frac{\partial^2 E}{\partial \theta_c^2} & 0 \\ 0 & 0 & \frac{\partial^2 E}{\partial \psi^2} \end{bmatrix}\right|_{\chi^B}. \tag{S40}$$

Thus, at $\psi^*_{cd}$, the Hessian has the eigenpair $\left(\left.\frac{\partial^2 E}{\partial \psi^2}\right|_{\chi^B}, [0,0,1]^T\right)$, indicating that near $\psi^*_{cd}$, the curvature of the energy as a function of orientation is given by $\left.\frac{\partial^2 E}{\partial \psi^2}\right|_{\chi^B}$. Consider then the fixed point of the system at this orientation, $\chi^*_{cd} = (a^*_{cd}, w^*_{cd}, \psi^*_{cd})$. The eigenvalue of the Hessian of the energy at this fixed point is

$$\begin{aligned}
\left.\frac{\partial^2 E}{\partial \psi^2}\right|_{\chi^*_{cd}} &= {a^*_{cd}}^2 \sum_{j \in K_{\text{act}}} \left(\sin(\theta_j - \psi^*_{cd}) - \frac{1}{N}\sum_{k \in K_{\text{act}}} W^{\text{sym}}_{jk} \sin(\theta_k - \psi^*_{cd})\right)\sin(\theta_j - \psi^*_{cd}) \\
&= {a^*_{cd}}^2 \sum_{j \in K_{\text{act}}} \left(\sin(\theta_j - \psi^*_{cd}) - \frac{1}{N}\sum_{k \in K_{\text{act}}} (J_I + J_E\cos(\theta_j - \theta_k))\sin(\theta_k - \psi^*_{cd})\right)\sin(\theta_j - \psi^*_{cd}) \\
&= {a^*_{cd}}^2 \sum_{j \in K_{\text{act}}} \Bigg(\sin(\theta_j - \psi^*_{cd}) - \frac{J_E}{N}\sum_{k \in K_{\text{act}}} \big(\cos(\theta_j - \psi^*_{cd})\cos(\theta_k - \psi^*_{cd}) \\
&\qquad + \sin(\theta_j - \psi^*_{cd})\sin(\theta_k - \psi^*_{cd})\big)\sin(\theta_k - \psi^*_{cd})\Bigg)\sin(\theta_j - \psi^*_{cd}) \\
&= {a^*_{cd}}^2 \sum_{j \in K_{\text{act}}} \sin^2(\theta_j - \psi^*_{cd})\left(1 - \frac{J_E}{N}\sum_{k \in K_{\text{act}}} \sin^2(\theta_k - \psi^*_{cd})\right).
\end{aligned} \tag{S41}$$

This indicates that the curvature of the energy around fixed points $\chi^*_{cd}$ is a function of the strength of local excitation in the system, $J_E$. The stability of these fixed points can be determined directly from the sign of (S41), and it can further be tuned between being stable and unstable by proper modulation of $J_E$. Most importantly, however, by setting

$$\frac{1}{J^*_E} = \frac{1}{N}\sum_{k \in K_{\text{act}}} \sin^2(\theta_k - \psi^*_{cd}), \tag{S42}$$

we can actually force the curvature as a function of orientation about $\chi^*_{cd}$ to zero, thereby generating a marginally stable fixed point about $\chi^*_{cd}$. We will denote these marginally stable fixed points by $\chi^{\text{ms}}(\psi) = (a^{\text{ms}}(\psi), w^{\text{ms}}(\psi), \psi)$, where $\psi \in (\psi^*_{dc} - \Delta\psi_{\text{ms}}/2, \psi^*_{dc} + \Delta\psi_{\text{ms}}/2)$, and where $\Delta\psi_{\text{ms}}$ is the width of the interval about $\psi^*_{cd}$ for which the marginally stable solution exists. We will refer to these values $J_E$ that generate marginally stable solutions as "optimal", and we will denote them by $J^*_E$.

#### Performance of Optimal Solutions

Here we analyze the marginally stable fixed point solutions that arise when $J_E = J^*_E$. Note that in (S42), $J^*_E$ is truly a function of $N$ and the number of "active" neurons $N_{\text{act}}$ (the size of the support of the bump, equal to the number of indices in $K_{\rm act}$), since $\psi^*_{cd}$ in (S29) can be entirely determined via $N$. Evaluating (S42) for different possible values of $N_{\rm act}$ shows that $J^*_E \in \Omega_{J_E}$ for $N_{\rm act} = 2, \ldots, N-2$, indicating that there are $N-3$ values of $J^*_E$ that generate marginally stable fixed point solutions. Considering $J^*_E$ as a function of $N_{\rm act}$ and $N$ alone, we can further simplify (S42) to

$$\frac{1}{J^*_{E,N_{\rm act}}} = \frac{1}{4} + \frac{1}{2N}\left(\tilde{n} + \frac{\sin(\tilde{n}\Delta\theta)}{\sin(\Delta\theta)}\right), \quad \tilde{n} = N_{\rm act} - \frac{N}{2}, \tag{S43}$$

highlighting that the inverse of these optimal $J^*_E$ symmetrically fill in the inverse parameter space ($\Omega^{-1}_{J_E} = \{1/J_E | J_E \in \Omega_{J_E}\}$) about $J^*_E = 4$ (corresponding to $N_{\rm act} = N/2$ for $N$ even) as $N$ increases (Fig 2d).

**Dynamics in the Absence of Input Velocity.** To characterize the performance of the marginally stable solutions in the absence of velocity input (i.e., $v_{\rm in} = 0$), we continue to use the energy landscape of the system. Specifically, we numerically calculate the eigenvalues and eigenvectors of the Hessian as a function of bump orientation. For each $\psi^B \in [0, 2\pi)$, we compute $w^B$ via (S30) and $a^B$ via (S31), and then we numerically calculate the eigenvalues and eigenvectors of $H(E)$ at $\chi^B = (a^B, w^B, \psi^B)$. Proceeding in this way, we find that the Hessian has a zero eigenvalue across all orientations, with corresponding eigenvectors pointing in directions that include a change in orientation (SI Fig S2). The existence of these zero eigenvalues, together with the fact that they appear when the Hessian is evaluated at $\chi^B$ that satisfies (S30)-(S31), indicates that the optimal values of local excitation $J^*_E$ not only locally flatten the energy landscape about $\chi^*_{cd}$, but also generate a flat basin across all orientations $\psi \in [0, 2\pi)$, with $a^{\rm ms}(\psi)$ and $w^{\rm ms}(\psi)$ varying appropriately to satisfy (S28),(S30)-(S31). Thus, $\Delta\psi_{\rm ms} = \Delta\theta$, and these values $J^*_E$ actually generate ring attractor solutions, allowing systems with as few as $N = 4$ neurons to accurately encode any orientation $\psi$ within the continuous interval $[0, 2\pi)$.

One striking feature of these optimal solutions (that is, the ring attractor solutions generated by optimal $J^*_E$) is that the number of active neurons $N_{\rm act}$ remains constant at each orientation $\psi \in [0, 2\pi)$. To maintain this, whenever an active neuron (or neuron in the support of the bump) becomes inactive (or leaves the support), a new neuron becomes active (or joins the support), thereby keeping $N_{\rm act}$ fixed as the orientation changes. This allows us to consider the subsystem of active neurons

$$\tau \dot{h}_j = -h_j + c_{ff} + \frac{1}{N}\sum_{k\in K_{\rm act}} W^{\rm sym}_{jk} h_k, \quad j \in K_{\rm act}, \tag{S44}$$

as a linear system. Taking $W^0_{N_{\rm act}}$ to be the $N_{\rm act} \times N_{\rm act}$ "active submatrix" along the diagonal of $W^0 = \frac{1}{\tau}(-I + W^{\rm sym}/N)$, we can rewrite (S44) as

$$\dot{\vec{h}}_{\rm act} = W^0_{N_{\rm act}} \vec{h}_{\rm act} + c_{ff}\vec{e}, \tag{S45}$$

where $\vec{h}_{\rm act} \in \mathbb{R}^{N_{\rm act}}_+$ denotes the vector of inputs to the active neurons, $\vec{e} \in \mathbb{R}^{N_{\rm act}}$ is the vector of all ones, $\vec{e} = [1, \ldots, 1]^T$, and we let the constant $c_{ff}$ absorb the inverse neural time constant $1/\tau$. We find the leading eigenvalue of $W^0_{N_{\rm act}}$ to be $0$ whenever $J_E = J^*_{E,N_{\rm act}}$, $N_{\rm act} = [2, \ldots, N-2]$ (SI Fig S4). This indicates that the "active" subsystem generates a line attractor. As a result, one can think of these optimal ring attractor solutions as being line attractors stitched together at each point where the support of the bump changes. Note that the presence of a zero leading eigenvalue in $W^0_{N_{\rm act}}$ when $J_E = J^*_E$ indicates that these "optimal" values of $J_E$ render the system degenerate. Although such degeneracies are unlikely to occur in biology (due to the strict tuning required), we find the degenerate system to be a useful tool in analyzing the "non-optimal" system, for which $J_E \neq J^*_E$.

**Dynamics in the Presence of Input Velocity.** When $v_{\rm in} \neq 0$, the optimal solutions again keep the number of active neurons approximately constant. As a result, we can once again work in the linear subsystem of active neurons:

$$\dot{\vec{h}}_{\rm act} = W^{v_{\rm in}}_{N_{\rm act}} \vec{h}_{\rm act} + c_{ff}\vec{e}, \tag{S46}$$

where now $W^{v_{\rm in}}_{N_{\rm act}}$ is the $N_{\rm act} \times N_{\rm act}$ "active submatrix" along the diagonal of $W^{v_{\rm in}} = \frac{1}{\tau}(-I + (W^{\rm sym} + v_{\rm in}W^{\rm asym})/N)$, and everything else is as above (note that (S45) is a special case of (S46), with $v_{\rm in} = 0$). Simulating the system for a range of constant $v_{\rm in}$ (where, without loss of generality, we choose $v_{\rm in} > 0$) suggests that the system will linearly integrate the input velocity $v_{\rm in}$ (including extremely small $v_{\rm in}$) as desired (see Fig 2f). This is not surprising given the stationary properties already determined in *Supplemental Information |Performance of Optimal Solutions |Dynamics in the Absence of Input Velocity*.

#### Performance of Non-Optimal Solutions

To dissect the performance of non-optimal solutions (that is, the solutions to systems with $J_E \neq J_E^*$), we will continue to use the linearity of the subsystem of active neurons. Unlike the optimal case, however, the number of active neurons $N_{\rm act}$ will vary not only as a function of the strength of local excitation $J_E$, but also as a function of orientation $\psi^B$ for any given $J_E \neq J_E^*$. As a result, the dynamics are governed by two different linear subsystems: one with $N_{\rm act} = n$ active neurons, and one with $N_{\rm act} = n+1$ active neurons. As found below, these will correspond to a stable and unstable system, respectively. The value of $n \in \{1, \ldots, N-2\}$ will depend on $J_E$ and can be determined by

$$n(J_E) = \begin{cases} N-2, & 2 < J_E < J_{E,N-2}^* \\ \tilde{n}, & J_{E,\tilde{n}+1}^* < J_E < J_{E,\tilde{n}}^*,\ \tilde{n} \in \{2, \ldots, N-3\} \\ 1, & J_{E,2}^* < J_E < \infty \end{cases}. \tag{S47}$$

We note that values of $J_E$ at either extreme (i.e., $J_E > J_{E,2}^*$ and $2 < J_E < J_{E,N-2}^*$) result in solutions that might be biologically implausible. At one end of this extreme, $J_E > J_{E,2}^*$, the system would have to encode a subset of orientations with only a single active neuron. At the other end, $2 < J_E < J_{E,N-2}^*$, the system would be required to keep $J_E$ tightly tuned in order to prevent the evolution to a homogeneous solution (when $J_E < 2$), since the interval $(2, J_{E,N-2}^*)$ is small and becomes increasingly so as network size $N$ increases. For this reason, we focus our analysis in the main text to intermediate values of $J_E \in [J_{E,N-2}^*, J_{E,2}^*]$. Mathematically, these analyses can be extended, and we treat the more general case ($J_E \in (2, \infty) = \Omega_{J_E}$) in this supplement.

Due to the rotational invariance of $W^{v_{\rm in}}$, $W^{v_{\rm in}}_{N_{\rm act}}$ will not change for fixed $N_{\rm act}$, regardless of which subset of contiguous subset of neurons is active. Hence, in what follows, we will analyze the system (S46) over a single angular unit of length $\Delta\theta$. The transition from $N_{\rm act} = n$ to $N_{\rm act} = n+1$ active neurons (or vice versa) will change the eigenvalues of $W^{v_{\rm in}}_{N_{\rm act}}$, thereby changing the underlying system dynamics and trajectory that the bump orientation $\psi^B$ traces out over time. Since the dynamics of the active neurons are governed by the linear system (S46), we expect the bump orientation in these two different regimes to be exponential in nature. We therefore take

$$\psi_s^B(t') = \alpha_s + \beta_s \exp(\lambda_s t'), \tag{S48}$$
$$\psi_u^B(t) = \alpha_u + \beta_u \exp(\lambda_u t) \tag{S49}$$

to represent the bump orientation in the stable and unstable regimes, respectively. In what follows, we will first analyze the drift of the bump orientation when $v_{\rm in} = 0$ to determine the drift rates $\lambda_s, \lambda_u$ and to identify the orientation $\psi_{\Delta n}$ at which the bump transitions between regimes. We then investigate the effect of small $v_{\rm in} \neq 0$ for which the values of $\lambda_s, \lambda_u, \psi_{\Delta n}$ are approximately unchanged (see *Methods |Model Analytics |Small Velocity Approximation*).

**Dynamics in the Absence of Input Velocity.** When $v_{\rm in} = 0$, the bump $\psi^B$ will drift away from unstable orientations and toward a stable one. In what follows, we consider the stable and unstable orientations, $\psi^s$ and $\psi^u$, within a single angular unit. Without loss of generality, we will assume $\psi^s < \psi^u$. Note that, for any non-optimal $J_E$, the stable and unstable orientations are given by (S29), with $c = d$ for one orientation, and $c = d+1$ for the other. Hence, we can take $\psi^u = \psi^s + \Delta\theta/2$.

When $\psi^B$ is sufficiently close to $\psi^s$, the system will be in the stable regime, and $\psi^B$ will be given by (S48). Further, we should have $\psi_s^B \to \psi^s$ as $t \to \infty$. Assuming $\lambda_s < 0$, we then have $\alpha_s = \psi^s$. Suppose in this regime there are $n_s$ active neurons. Then, from (S27) we have

$$\tau \beta_s \lambda_s e^{t\lambda_s} = J_E f_{\rm odd}(w^B, \psi^s + \beta_s e^{t\lambda_s}). \tag{S50}$$

Consider

$$
\begin{aligned}
f_{\text{odd}}(w^B, \psi^s + \beta_s e^{t\lambda_s}) &= \frac{1}{N} \sum_{k\in K_{\text{act}}} \Big( \cos\big(\theta_k - (\psi^s + \beta_s e^{\lambda_s t})\big) - \cos(w^B/2) \Big) \sin\big(\theta_k - (\psi^s + \beta_s e^{\lambda_s t})\big) \\
&= \frac{1}{N} \sum_{k\in K_{\text{act}}} \big( \cos(\theta_k - \psi^s)\cos(\beta_s e^{\lambda_s t}) + \sin(\theta_k - \psi^s)\sin(\beta_s e^{\lambda_s t}) - \cos(w^B/2) \big) \\
&\qquad\qquad \big( \sin(\theta_k - \psi^s)\cos(\beta_s e^{\lambda_s t}) - \cos(\theta_k - \psi^s)\sin(\beta_s e^{\lambda_s t}) \big).
\end{aligned}
$$

Since $\psi^s$ is given by (S29), the summations over odd functions centered at $\psi^s$ will cancel themselves out (being anti-symmetric about $\psi^s$). This gives us

$$
\begin{aligned}
f_{\text{odd}}(w^B, \psi^s + \beta_s e^{t\lambda_s}) = -\frac{\sin(\beta_s e^{\lambda_s t})}{N} \sum_{k\in K_{\text{act}}} \Big[ \big( \cos(\beta_s e^{\lambda_s t})\cos(\theta_k - \psi^s) - \cos(w^B/2) \big) \cos(\theta_k - \psi^s) \\
- \cos(\beta_s e^{\lambda_s t}) \sin^2(\theta_k - \psi^s) \Big].
\end{aligned}
$$

Taking $t$ large, we use small angle approximations to get

$$
f_{\text{odd}}(w^B, \psi^s + \beta_s e^{t\lambda_s}) \approx \beta_s e^{\lambda_s t} \left( \frac{1}{J^*_{E,n_s}} - f_{\text{even}}(w^B, \psi^s) \right).
$$

Assuming $w^B \approx w^s$, where $\chi^s = (a^s, w^s, \psi^s)$ is the stable fixed point associated with $\psi^s$, and using (S30) and (S50), we find

$$
\lambda_s \approx \frac{1}{\tau} \left( \frac{J_E}{J^*_{E,n_s}} - 1 \right). \tag{S51}
$$

Thus, the drift rate $\lambda_s$ will depend on how close $J_E$ is to the optimal value $J^*_{E,n_s}$, with closer $J_E$ corresponding to slower drift. Note that our assumption that $\lambda_s < 0$ requires

$$
J_E < J^*_{E,n_s}. \tag{S52}
$$

Similarly, when $\psi^B$ is sufficiently close to $\psi^u$, the system will be in the unstable regime, and $\psi^B$ will be given by (S49). If we now take $t \to -\infty$, we should have $\psi^B_u \to \psi^u$. Assuming now $\lambda_u > 0$, this indicates that $\alpha_u = \psi^u$. Using the same types of approximations as above, we find

$$
\lambda_u \approx \frac{1}{\tau} \left( \frac{J_E}{J^*_{E,n_u}} - 1 \right). \tag{S53}
$$

where $n_u$ is the number of active neurons. The assumptions that $\lambda_u > 0$ requires

$$
J_E > J^*_{E,n_u}. \tag{S54}
$$

Since $J^*_{E,N-2} < J^*_{E,N-3} < \cdots < J^*_{E,2}$, inequalities (S52) and (S54) imply that $n_u = n_s + 1$. Hence, $n(J_E) = n_s$, as defined above in (S47). Note that, since $\lambda_u$ was found by evolving the linear dynamics in negative time, this corresponds to the "repulsion" rate. That is, $\lambda_u$ indicates the rate at which $\psi^B$ will be pushed away from the unstable orientation $\psi^u$. Again, we find from (S53) that this rate will depend on how close $J_E$ is to $J^*_{E,n+1}$, being smaller the closer $J_E$ is to this optimal value.

Given the strictly decreasing nature of the optimal $J^*_{E,n}$ as a function of $n$, and given the relations (S52), (S54) found between $J_E$ and $J^*_E$ when $\psi^B$ is moving towards a stable orientation $\psi^s$ or away from an unstable orientation $\psi^u$, respectively, the linear active subsystem will be stable when $N_{\text{act}} = n$ and unstable when $N_{\text{act}} = n + 1$. Numerically computing the leading eigenvalue of $W^0_n$ and $W^0_{n+1}$ confirms this. Further, we find our approximations of these leading eigenvalues, $\lambda_s$ and $\lambda_u$, to be close fits to the numerically computed values (Fig S4).

To understand how much these stable and unstable dynamics affect the overall trajectories of the bump, we next calculate the orientation $\psi_{\Delta n}$ at which the system will transition between these stable and unstable regimes. To find this transition point, we again consider the stable and unstable orientations, $\psi^s$ and $\psi^u = \psi^s + \Delta\theta/2$, respectively, within a single angular unit with length $\Delta\theta$. We assume the system starts in the unstable regime at some orientation $\psi_u^B(0) = \psi_0 < \psi^u$, implying $\beta_u = \psi_0 - \psi^u$. We define $t = t_{\Delta n}$ to be the time when the orientation reaches this transition point from the unstable regime, and we define $t' = 0$ to be the time when the system crosses into the stable regime. This gives

$$\psi_u^B(t_{\Delta n}) = \psi_{\Delta n}, \tag{S55}$$
$$\psi_s^B(0) = \psi_{\Delta n}, \tag{S56}$$

where $\psi_u^B$, $\psi_s^B$ are given by (S49), (S48), respectively, with $\alpha_u = \psi^u = \psi^s + \Delta\theta/2$ and $\alpha_s = \psi^s$. Further, we assume this transition is smooth, requiring

$$\dot{\psi}_u^B(t_{\Delta n}) = \dot{\psi}_s^B(0). \tag{S57}$$

Condition (S56) implies that $\beta_s = \psi_{\Delta n} - \psi^s$. We can now rewrite (S48)-(S49) as

$$\psi_s^B(t') = \psi^s + (\psi_{\Delta n} - \psi^s)\exp(-|\lambda_s|t'), \tag{S58}$$
$$\psi_u^B(t) = \psi^u + (\psi_0 - \psi^u)\exp(\lambda_u t). \tag{S59}$$

Conditions (S55) and (S57) imply that:

$$\begin{aligned}
\psi_u^B(t_{\Delta n}) = \psi_{\Delta n} &\implies \psi_{\Delta n} = \psi^s + \frac{\Delta\theta}{2} + \beta_u \exp(\lambda_u t_{\Delta n}) \\
&\implies \beta_u \exp(\lambda_u t_{\Delta n}) = \psi_{\Delta n} - \left(\psi^s + \frac{\Delta\theta}{2}\right), \\
\dot{\psi}_u^B(t_{\Delta n}) = \dot{\psi}_s^B(0) &\implies \beta_u \lambda_u \exp(\lambda_u t_{\Delta n}) = -(\psi_{\Delta n} - \psi^s)|\lambda_s| \\
&\implies \lambda_u \left(\psi_{\Delta n} - \left(\psi^s + \frac{\Delta\theta}{2}\right)\right) = -(\psi_{\Delta n} - \psi^s)|\lambda_s| \\
&\implies \psi_{\Delta n} = \psi^s + \frac{\Delta\theta}{2}\left(\frac{1}{1 + |\lambda_s|/\lambda_u}\right).
\end{aligned} \tag{S60}$$

The stable regime spans a width $\psi_{\Delta n} - \psi^s$ on either side of the stable fixed point $\psi^s$, and thus the total width $\Delta\theta_s$ of the stable regime is:

$$\Delta\theta_s = 2(\psi_{\Delta n} - \psi^s) = \Delta\theta\left(\frac{1}{1 + |\lambda_s|/\lambda_u}\right). \tag{S61}$$

The width $\Delta\theta_u$ of the unstable regime is then:

$$\Delta\theta_u = \Delta\theta - \Delta\theta_s = \Delta\theta\left(\frac{1}{1 + \lambda_u/|\lambda_s|}\right). \tag{S62}$$

The bump will cross from the unstable regime into the stable regime at a time $t_{\Delta n}$ that satisfies $\psi_u^B(t_{\Delta n}) = \psi_{\Delta n}$. Solving (S59) for $t_{\Delta n}$ yields

$$t_{\Delta n} = \frac{1}{\lambda_u}\log\left(\frac{\Delta\theta_u}{2(\psi^u - \psi_0)}\right). \tag{S63}$$

Equations (S58)-(S59) can similarly be used to determine the total drift time $\tau_d$ that it takes for the bump to drift from within $\varepsilon_u$ of an unstable fixed point to within $\varepsilon_s$ of a stable fixed point. This is given by the time it takes for the bump to cross into the stable regime (given by (S63) with $\psi_0 = \psi^u(0) - \varepsilon_u$), plus the additional time $t_s$ that it takes for the bump to cross the stable regime, where $t_s$ satisfies $\psi_s^B(t_s) = \psi^s(0) + \varepsilon_s$. Solving (S58) for $t_s$ yields a total drift time of:

$$\tau_d = \frac{1}{|\lambda_u|}\log\left(\frac{\Delta\theta_u}{2\varepsilon_u}\right) + \frac{1}{|\lambda_s|}\log\left(\frac{\Delta\theta_s}{2\varepsilon_s}\right). \tag{S64}$$

Taking $\varepsilon_u = \Delta\theta_u/2e$ and $\varepsilon_s = \Delta\theta_s/2e$, (S64) reduces to:

$$\tau_d = \frac{1}{|\lambda_u|} + \frac{1}{|\lambda_s|}. \tag{S65}$$

Note that this corresponds to the time it takes the bump to travel an angular distance $\Delta\psi_d = (1 - 1/e)\Delta\theta/2$. In the main text, we used $\Delta\psi_d/\tau_d$ as a measure of the net drift speed, which can be expressed as

$$|\lambda_d| = \frac{\Delta\psi_d}{\tau_d} = \left(1 - \frac{1}{e}\right)\frac{\Delta\theta}{2}\frac{|\lambda_u||\lambda_s|}{|\lambda_u| + |\lambda_s|} \tag{S66}$$

$$= c\Delta\theta_s|\lambda_s| \tag{S67}$$

$$= c\Delta\theta_u|\lambda_u|, \tag{S68}$$

where $c = (e-1)/2e$ is a constant.

**Dynamics in the Presence of Small Input Velocity** In the presence of small velocity input $v_{\text{in}}$, the stable and unstable drift rates remain approximately unchanged (see *Methods|Model Analytics|Small Velocity Approximation*), and thus so too does the location $\psi_{\Delta n}$ of the boundary between stable and unstable regimes (see (S60)). However, the locations of the stable and unstable orientations will change with $v_{\text{in}}$. That is, $\psi^s = \psi^s(v_{\text{in}})$ and $\psi^u = \psi^u(v_{\text{in}})$. The stable and unstable fixed point orientations referenced above and given by (S29) are then $\psi^s(0)$ and $\psi^u(0)$, respectively. To determine the new orientations of these fixed points, we consider a bump that begins at the stable orientation $\psi^s(0)$ with an initial bump velocity $v_{\text{in}}$. Without loss of generality, we assume that $v_{\text{in}} > 0$ and thus drives the bump toward $\psi^u(0) = \psi^s(0) + \Delta\theta/2$. The bump orientation will again be governed by (S48)-(S49), but with a modified set of initial conditions:

$$\dot{\psi}_s^B(0) = v_{\text{in}}, \tag{S69}$$

$$\psi_s^B(0) = \psi^s(0), \tag{S70}$$

$$\dot{\psi}_s^B(t_{\Delta n}) = \dot{\psi}_u^B(0), \tag{S71}$$

$$\psi_s^B(t_{\Delta n}) = \psi_u^B(0) = \psi_{\Delta n}, \tag{S72}$$

where $t' = t_{\Delta n}$ and $t = 0$ are now the times that $\psi_s^B = \psi_{\Delta n}$ and $\psi_u^B = \psi_{\Delta n}$, respectively. Initial condition (S69) implies that $\beta_s = -v_{\text{in}}/|\lambda_s|$, while initial condition (S70) implies that $\alpha_s = \psi^s(0) + v_{\text{in}}/|\lambda_s|$. Initial conditions (S71)-(S72) further imply that:

$$\begin{aligned}
\dot{\psi}_s^B(t_{\Delta n}) = \dot{\psi}_u^B(0) &\implies v_{\text{in}}\exp(-|\lambda_s|t_{\Delta n}) = \lambda_u\beta_u, \\
\psi_s^B(t_{\Delta n}) = \psi_{\Delta n} &\implies \psi^s(0) + \frac{v_{\text{in}}}{|\lambda_s|}\Big(1 - \exp(-|\lambda_s|t_{\Delta n})\Big) = \psi_{\Delta n} \\
&\implies \psi^s(0) + \frac{v_{\text{in}}}{|\lambda_s|}\left(1 - \frac{\lambda_u\beta_u}{v_{\text{in}}}\right) = \psi_{\Delta n} \\
&\implies \beta_u = \frac{v_{\text{in}}}{\lambda_u} - (\psi_{\Delta n} - \psi^s(0))\frac{|\lambda_s|}{\lambda_u} \\
&\implies \beta_u = \frac{v_{\text{in}}}{\lambda_u} - \left(\frac{\Delta\theta}{2} - (\psi_{\Delta n} - \psi^s(0))\right) \\
&\implies \beta_u = \frac{v_{\text{in}}}{\lambda_u} - (\psi^u(0) - \psi_{\Delta n}), && \text{(S73)} \\
\psi_u^B(0) = \psi_{\Delta n} &\implies \alpha_u + \beta_u = \psi_{\Delta n} \\
&\implies \alpha_u = \psi_{\Delta n}\left(1 + \frac{|\lambda_s|}{\lambda_u}\right) - \psi^s(0)\frac{|\lambda_s|}{\lambda_u} - \frac{v_{\text{in}}}{\lambda_u} \\
&\implies \alpha_u = \psi^s(0) + \frac{\Delta\theta}{2} - \frac{v_{\text{in}}}{\lambda_u} \\
&\implies \alpha_u = \psi^u(0) - \frac{v_{\text{in}}}{\lambda_u}. && \text{(S74)}
\end{aligned}$$

Together, this yields:

$$\psi_s^B(t') = \psi^s(0) + \frac{v_{\text{in}}}{|\lambda_s|}\Big(1 - \exp(-|\lambda_s|t')\Big), \tag{S75}$$

$$\psi_u^B(t) = \psi^u(0) - \frac{v_{\text{in}}}{\lambda_u} + \left(\frac{v_{\text{in}}}{\lambda_u} - (\psi^u(0) - \psi_{\Delta n})\right)\exp(\lambda_u t). \tag{S76}$$

The bump will cross from the stable to the unstable regimes at a time $t_{\Delta n}$, where

$$t_{\Delta n} = \frac{1}{|\lambda_s|}\log\left(\frac{1}{1 - |\lambda_s|\Delta\theta_s/(2v_{\text{in}})}\right) \tag{S77}$$

can be determined from (S75).

In the limit that $t \to -\infty$, the bump will be driven toward (and hence, in forward time, away from) an unstable fixed point with orientation

$$\psi^u(v_{\text{in}}) = \psi^u(0) - v_{\text{in}}/\lambda_u. \tag{S78}$$

Similarly, in the limit that $t' \to \infty$, the bump will be driven toward a stable fixed point with orientation

$$\psi^s(v_{\text{in}}) = \psi^s(0) + v_{\text{in}}/|\lambda_s|. \tag{S79}$$

For $v_{\text{in}} = 0$, the stable and unstable fixed point orientations are located at $\psi^s = \psi^s(0)$ and $\psi^u = \psi^u(0)$ given by (S29), as above. As $v_{\text{in}}$ increases, both fixed point orientations shift toward the boundary between the stable and unstable regimes at $\psi_{\Delta n}$ (the stable orientation shifts in the same direction as $v_{\text{in}}$, while the unstable orientation shifts in the opposite direction as $v_{\text{in}}$). The stable fixed point orientation will reach this boundary at a threshold velocity $v_{\text{thresh}}$ given by:

$$\begin{aligned} v_{\text{thresh}} &= |\lambda_s|(\psi_{\Delta n} - \psi^s(0)) \\ &= |\lambda_d|/2c, \end{aligned} \tag{S80}$$

where $c = (e-1)/2e$, as above, and where we have used (S67) to express the threshold velocity in terms of the net drift speed $|\lambda_d|$. At the threshold velocity, the unstable fixed point orientation will have also shifted to this same boundary:

$$\begin{aligned} \psi^u(v_{\text{thresh}}) &= \psi^u(0) - v_{\text{thresh}}/\lambda_u \\ &= \psi^s(0) + \frac{\Delta\theta}{2} - \frac{|\lambda_s|}{\lambda_u}(\psi_{\Delta n} - \psi^s(0)) \\ &= \psi^s(0) + \frac{1}{2}(\Delta\theta - \Delta\theta_u) \\ &= \psi_{\Delta n}. \end{aligned}$$

Above this threshold velocity, the location of the stable fixed point orientation is beyond the boundary of the stable regime. As a result, the bump will be pulled toward this orientation but will never reach it; instead, the bump will cross the boundary into the unstable regime, where it will be pushed away from an unstable fixed point. The orientation of this unstable fixed point is similarly beyond the boundary of the unstable regime, and pushes the bump from behind. As a result, the bump will be pulled toward and pushed away from stable and unstable orientations that it can't reach. This will cause the bump to decelerate and accelerate as it travels through the stable and unstable regimes, respectively. The bump will be moving the slowest as it passes the boundary from the stable into the unstable regime (i.e., at $\psi_s^B = \psi_{\Delta n}$); beyond this point, it will begin accelerating away from the unstable fixed point. To find the velocity at this point, note that

$$\psi_s^B(t_{\Delta n}) = \psi_{\Delta n} \implies \psi_{\Delta n} = \psi^s(0) + \frac{v_{\text{in}}}{|\lambda_s|}(1 - \exp(-|\lambda_s| t_{\Delta n}))$$
$$\implies \exp(-|\lambda_s| t_{\Delta n}) = 1 - (\psi_{\Delta n} - \psi^s(0))\frac{|\lambda_s|}{v_{\text{in}}}.$$

The slowest bump velocity $\nu_{\min}$ is thus given by

$$\begin{aligned}\nu_{\min} = \dot{\psi}_s^B(t_{\Delta n}) &= v_{\text{in}} \exp(-|\lambda_s| t_{\Delta n}) \\ &= v_{\text{in}}\left(1 - (\psi_{\Delta n} - \psi^s(0))\frac{|\lambda_s|}{v_{\text{in}}}\right) \\ &= v_{\text{in}} - |\lambda_s|(\psi_{\Delta n} - \psi^s(0)) \\ &= v_{\text{in}} - v_{\text{thresh}}.\end{aligned} \tag{S81}$$

The bump will be moving the fastest as it passes the boundary from the unstable into the stable regime (i.e., at $\psi_u^B = \psi_{\Delta n} + \Delta\theta_u = 2\psi^u(0) - \psi_{\Delta n}$); beyond this point, it will begin decelerating toward the stable fixed point. To find the velocity at this point, note that

$$\psi_u^B(t_{\Delta n}) = 2\psi^u(0) - \psi_{\Delta n} + \implies 2\psi^u(0) - \psi_{\Delta n} = \psi^u(0) - \frac{v_{\text{in}}}{\lambda_u} + \left(\frac{v_{\text{in}}}{\lambda_u} - \big(\psi^u(0) - \psi_{\Delta n}\big)\right)\exp(\lambda_u t_{\Delta n})$$
$$\implies \left(\frac{v_{\text{in}}}{\lambda_u} - \big(\psi^u(0) - \psi_{\Delta n}\big)\right)\exp(\lambda_u t_{\Delta n}) = \frac{v_{\text{in}}}{\lambda_u} - \big(\psi_{\Delta n} - \psi^u(0)\big).$$

The fastest bump velocity $\nu_{\max}$ will thus be given by

$$\begin{aligned}\nu_{\max} = \dot{\psi}_u^B(t_{\Delta n}) &= \lambda_u\left(\frac{v_{\text{in}}}{\lambda_u} - \big(\psi^u(0) - \psi_{\Delta n}\big)\right)\exp(\lambda_u t_{\Delta n}) \\ &= \lambda_u\left(\frac{v_{\text{in}}}{\lambda_u} - \big(\psi_{\Delta n} - \psi^u(0)\big)\right) \\ &= \lambda_u\left(\frac{v_{\text{in}}}{\lambda_u} + \frac{|\lambda_s|}{\lambda_u}\big(\psi_{\Delta n} - \psi^s(0)\big)\right) \\ &= v_{\text{in}} + v_{\text{thresh}}.\end{aligned} \tag{S82}$$

In the main text, we define the linearity of integration $\ell$ to be

$$\ell = \frac{\nu_{\min}}{\nu_{\max}} = \frac{v_{\text{in}} - v_{\text{thresh}}}{v_{\text{in}} + v_{\text{thresh}}}. \tag{S83}$$

**Degradation of Performance as a Function of Local Excitation.** In the main text, we consider three different measures of performance: the net drift speed, the threshold input velocity needed to move the bump continuously, and the linearity of integration for input velocities above the threshold value. To characterize how each of these performance measures locally degrades as we tune the local excitation away from an optimal value, we consider the derivative of each performance measure $P$ with respect to $J_E$ when evaluated at an optimal value of local excitation $J_E^*$:

$$m_P(J_E^*) = \left.\frac{\partial P}{\partial J_E}\right|_{J_E^*}, \tag{S84}$$

where $P \in \{|\lambda_d|, v_{\text{thresh}}, \ell\}$ is one of the net drift speed, threshold velocity, or linearity of integration, respectively. Because each performance measure can be written in terms of the net drift speed, we can rewrite this as

$$m_P(J_E^*) = \left.\frac{\partial P}{\partial |\lambda_d|}\frac{\partial |\lambda_d|}{\partial J_E}\right|_{J_E^*}, \tag{S85}$$

where

$$\frac{\partial v_{\text{thresh}}}{\partial|\lambda_d|} = \frac{1}{2c}, \tag{S86}$$

$$\begin{aligned}\frac{\partial \ell}{\partial|\lambda_d|} &= \frac{-2v_{\text{in}}}{(v_{\text{in}}+v_{\text{thresh}})^2}\frac{\partial v_{\text{thresh}}}{\partial|\lambda_d|} \\ &= \frac{-v_{\text{in}}}{c(v_{\text{in}}+v_{\text{thresh}})^2}.\end{aligned} \tag{S87}$$

In what follows, we first determine $\partial|\lambda_d|/\partial J_E|_{J_E^*}$, and we then use (S85)-(S87) to determine the rate at which performance degrades around the optimal values $J_E^*$. Taking the derivative of (S66) with respect to $J_E$, we find

$$\begin{aligned}\frac{\partial|\lambda_d|}{\partial J_E} &= c\Delta\theta\left(\frac{(|\lambda_s|+|\lambda_u|)(|\lambda_s|'+|\lambda_u|')-|\lambda_s||\lambda_u|(|\lambda_s|'+|\lambda_u|')}{(|\lambda_s|+|\lambda_u|)^2}\right) \\ &= c\Delta\theta\left(\frac{|\lambda_u|^2|\lambda_s|'+|\lambda_s|^2|\lambda_u|'}{(|\lambda_s|+|\lambda_u|)^2}\right),\end{aligned} \tag{S88}$$

where $|\lambda|' = \partial|\lambda|/\partial J_E$. From (S51), (S53), we have

$$|\lambda_u|+|\lambda_s| = \frac{J_E}{\tau}\left(\frac{J_{E,n}^*-J_{E,n+1}^*}{J_{E,n}^*J_{E,n+1}^*}\right). \tag{S89}$$

Recall from (S52) and (S54), we have $J_{E,n+1}^* < J_E < J_{E,n}^*$, such that (S89) is positive. Further,

$$|\lambda_s|' = \frac{-1}{\tau J_{E,n}^*}, \tag{S90}$$

$$|\lambda_u|' = \frac{1}{\tau J_{E,n+1}^*}, \tag{S91}$$

such that

$$|\lambda_u|^2|\lambda_s|'+|\lambda_s|^2|\lambda_u|' = \frac{1}{\tau^3}\left(\frac{J_{E,n}^*-J_{E,n+1}^*}{J_{E,n}^*J_{E,n+1}^*}\right)\left(1-\frac{J_E^2}{J_{E,n}^*J_{E,n+1}^*}\right). \tag{S92}$$

From (S89) and (S92), we find

$$\frac{\partial|\lambda_d|}{\partial J_E} = \frac{c\Delta\theta}{\tau}\left(\frac{J_{E,n}^*J_{E,n+1}^*}{J_{E,n}^*-J_{E,n+1}^*}\right)\left(\frac{1}{J_E^2}-\frac{1}{J_{E,n}^*J_{E,n+1}^*}\right). \tag{S93}$$

Letting $J_E$ approach some optimal value $J_E^*$ from below and above, respectively, we find

$$\lim_{J_E\nearrow J_E^*}\left.\frac{\partial|\lambda_d|}{\partial J_E}\right|_{J_E} = -\frac{c\Delta\theta}{\tau J_E^*}, \tag{S94}$$

$$\lim_{J_E\searrow J_E^*}\left.\frac{\partial|\lambda_d|}{\partial J_E}\right|_{J_E} = \frac{c\Delta\theta}{\tau J_E^*}. \tag{S95}$$

Note that (S94)-(S95) indicate that (S85) will change signs as $J_E$ passes through an optimal value (which is to be expected since the optimal values should correspond to local extrema of our performance measures), but will have a constant magnitude locally about the optimal $J_E^*$. Thus, for each performance measure $P$, the magnitude of the slope $|m_P(J_E^*)|$ gives us a local approximation for how performance changes as $J_E$ is tuned away from $J_E^*$. We can then estimate $P = P(J_E)$ around $J_E^*$ by

$$P(J_E) \approx (J_E - J_E^*)m_P(J_E^*) + P^*, \tag{S96}$$

where $P^* = P(J_E^*)$ indicates the best achievable performance ($\approx 0$ for $P = |\lambda_d|, v_{\text{thresh}}$, and $\approx 1$ for $P = \ell$). Let $P_{\text{desired}} = P^* + \varepsilon_P^{\text{tol}}$ indicate the desired performance threshold. Then, for sufficiently small $|\varepsilon_P^{\text{tol}}|$, the tolerance of the system about $J_E^*$ (defined to be the length of the interval containing $J_E^*$ that satisfies the desired performance threshold) is given approximately by

$$\text{tol}_P(J_E^*) \approx \frac{2\varepsilon_P^{\text{tol}}}{|m_P(J_E^*)|}. \tag{S97}$$

Combining (S94)-(S95) with (S86)-(S87), we have

$$m_{|\lambda_d|}(J_E^*) = \mp\frac{2\pi c}{\tau N J_E^*}, \tag{S98}$$

$$m_{v_{\text{thresh}}}(J_E^*) = \mp\frac{\pi}{\tau N J_E^*}, \tag{S99}$$

$$m_{\ell}(J_E^*) = \pm\frac{2\pi}{\tau v_{\text{in}} N J_E^*}, \tag{S100}$$

as $J_E$ approaches $J_E^*$ from below and above, respectively. Note that for $P = |\lambda_d|$ or $P = v_{\text{thresh}}$, $m_P(J_E^*)$ decreases away from the optimal value $J_E^*$, while for $P = \ell$, it increases. Since $|\lambda_d|$ and $v_{\text{thresh}}$ are performance measures that should be minimized and $\ell$ should be maximized (reaching a global maximum at $1$), this indicates that, for a desired performance threshold $P_{\text{desired}}$, our estimate will give a lower bound on the actual tolerance about $J_E^*$:

$$\text{tol}_P(J_E^*) \geq \frac{2\varepsilon_P^{\text{tol}}}{|m_P(J_E^*)|}. \tag{S101}$$

Thus, for each performance measure $P$, the resulting tolerance about $J_E^*$ is proportional to $1/|m_P(J_E^*)|$, which yields

$$\text{net drift speed: } \text{tol}_{|\lambda_d|}(J_E^*) \geq c_{|\lambda_d|} N J_E^*, \tag{S102}$$

$$\text{threshold velocity: } \text{tol}_{v_{\text{thresh}}}(J_E^*) \geq c_{v_{\text{thresh}}} N J_E^*, \tag{S103}$$

$$\text{linearity of integration: } \text{tol}_{\ell}(J_E^*) \geq c_{\ell} N J_E^* v_{\text{in}}, \tag{S104}$$

where

$$c_{|\lambda_d|} = \frac{\tau}{\pi c}\varepsilon_{|\lambda_d|}^{\text{tol}}, \tag{S105}$$

$$c_{v_{\text{thresh}}} = \frac{2\tau}{\pi}\varepsilon_{v_{\text{thresh}}}^{\text{tol}}, \tag{S106}$$

$$c_{\ell} = \frac{\tau v_{\text{in}}}{\pi}\varepsilon_{\ell}^{\text{tol}}. \tag{S107}$$

For a given network size $N$, we can compute the net volume $V$ of parameter space that satisfies a given performance threshold by summing the tolerances about each optimal values of local excitation:

$$\begin{aligned} V_P &\geq c_P N \sum_{N_{\text{act}}=2}^{N-2} J_{E,N_{\text{act}}}^* \\ &\geq 2N^2 \sum_{N_{\text{act}}=2}^{N-2} \frac{1}{N_{\text{act}} - \sin(2\pi N_{\text{act}}/N)/\sin(2\pi/N)}, \end{aligned} \tag{S108}$$

where $P \in \{|\lambda_d|, v_{\text{thresh}}, \ell\}$ is the given performance measure, and where we have used (S43) to rewrite $J_{E,N_{\text{act}}}^*$. The largest contribution to this sum will be for $N_{\text{act}} = 2$ (note that this is the only contribution for $N = 4$), which allows us to simplify the lower bound on the net volume to

$$V_P \geq c_P \frac{N^2}{1 - \cos(2\pi/N)}. \tag{S109}$$
